# RET activation controlled by MAB21L4-CacyBP interaction drives squamous cell carcinoma

**DOI:** 10.1101/2022.01.19.476979

**Authors:** Ankit Srivastava, Cristina Tommasi, Dane Sessions, Angela Mah, Tomas Bencomo, Jasmine M. Garcia, Tiffany Jiang, Michael Lee, Joseph Y. Shen, Lek Wei Seow, Audrey Nguyen, Kimal Rajapakshe, Cristian Coarfa, Kenneth Y. Tsai, Vanessa Lopez-Pajares, Carolyn S. Lee

## Abstract

Epithelial squamous cell carcinomas (SCC) most commonly originate in the skin, where they display disruptions in the normally tightly regulated homeostatic balance between keratinocyte proliferation and terminal differentiation. We performed a transcriptome-wide screen for genes of unknown function that possess inverse expression patterns in differentiating keratinocytes compared to cutaneous SCC (cSCC) and identified *MAB21L4* (*C2ORF54*) as an enforcer of terminal differentiation that suppresses carcinogenesis. Loss of MAB21L4 in human cSCC organoids enabled malignant transformation through increased expression of the receptor tyrosine kinase rearranged during transfection (RET). In addition to transcriptional upregulation of RET, MAB21L4 deletion preempted recruitment of the CacyBP-Siah1 E3 ligase complex to RET and reduced its ubiquitylation. Both genetic disruption of *RET* or selective RET inhibition with BLU-667 (pralsetinib) suppressed tumorigenesis in SCC organoids and in vivo tumors while inducing concomitant differentiation. Our results suggest that targeting RET activation is a potential therapeutic strategy for treating SCC.

**Statement of Significance:** Few targeted therapies are available to individuals with cSCC who seek or require non-surgical management. Our study demonstrates that downregulation of RET is required for epithelial differentiation and opposes carcinogenesis in cSCC as well as SCC arising from other epithelial tissues.

## Introduction

Squamous cell carcinomas (SCC) arising from epithelial tissues most commonly originate in the skin. PD-1 blockade with response rates of 35-50% remains the sole FDA-approved targeted approach for cutaneous SCC (cSCC) and is limited to patients with locally advanced or metastatic disease (1, 2). Curative surgery associated with rare instances of functional impairment is the standard of care for small tumors and may not be tolerated by all individuals. Additional targeted agents to treat cSCC, particularly those effective against early lesions, would therefore address this potential treatment gap by providing a non-invasive alternative to surgical management.

Loss of self-renewal and terminal differentiation are normally coupled in the epidermis, with keratinocytes dividing at controlled rates in the innermost basal layer and losing this ability as they differentiate and migrate outwards to the cornified layer (3). This regulation is notably absent in cSCC, and epidermal expression of oncogenic Ras in particular is well known to drive proliferation and transformation while inhibiting differentiation (4, 5). Impaired differentiation in cSCC is also suggested by whole-exome sequencing analyses that indicate 18% of tumors contain an inactivating mutation in *NOTCH1/2* (6). Genetic loss-of-function studies further support critical roles in tumorigenesis for differentiation-associated proteins, including loricrin and receptor-interacting protein kinase 4 (RIPK4) (7, 8). Similar to other cancers, poor tumor differentiation in cSCC predicts recurrence, metastasis, and worse overall survival (9, 10); however, the full suite of differentiation regulators disrupted in cSCC and their contributions to tumorigenesis have not been fully defined.

Uncharacterized open reading frames (ORFs) in the human genome frequently lack detectable homology to proteins of known function, which may delay their experimental characterization. Despite these challenges, an increasing number of genes originally from this pool are currently appreciated to control a diverse range of fundamental cellular and biological processes, including regulation of Wnt signaling (11), cell cycle progression (12), and effector T-cell homing (13). In our study, we searched this group of uncharacterized candidates for a regulator of epidermal differentiation and hypothesized that disrupting its function would drive cSCC carcinogenesis as well as uncover potential therapeutic targets. Through transcriptomic analyses comparing human cSCC to differentiating primary keratinocytes followed by functional interrogation in human skin organoids and proximity proteomics, we reveal a pro-differentiation, tumor-suppressive role for MAB21L4 (C2orf54) and its binding partner, CacyBP. Increased expression of the receptor tyrosine kinase (RTK) RET accompanies cSCC-perturbed MAB21L4-CacyBP interaction and exposes a vulnerability for which selective inhibitors are currently available. Both genetic ablation of *RET* or inhibition by pralsetinib (BLU-667) disrupt tumorigenesis in human SCC organoids and in vivo tumors while triggering concomitant differentiation, establishing initial proof of concept for targeting RET in SCC.

## Results

### Transcriptomics and proximity proteomics identify MAB21L4 and CacyBP as regulators of epidermal differentiation

Tumorigenesis is consistently accompanied by deregulated expression of epidermal differentiation mediators, although the complete range of affected genes and the nature of their functional roles have not been fully defined. To identify uncharacterized regulators of epidermal differentiation disrupted in malignancy, we interrogated RNA sequencing data from progenitor and differentiating human keratinocytes (14) as well as eight cSCC with patient-matched normal skin to complement three tumor-normal pairs previously reported (15) and focused on genes of unknown function. *MAB21L4* (*C2ORF54*), which encodes a 447 amino acid protein, was increased >480-fold in cultured primary keratinocytes on day 6 of differentiation in calcium-containing medium and significantly downregulated in human cSCC (**Fig. 1A-B**). Progressive induction of *MAB21L4* during keratinocyte differentiation was also confirmed at the protein level (**Fig. 1C**). Consistent with this pattern of expression, MAB21L4 protein was detected in the suprabasal compartment of normal human skin by immunofluorescence (**Supplementary Fig. S1A**). Subsequent analysis of publicly available microarray data for genes that possess Boolean implications with *MAB21L4* (16) revealed low expression of *MAB21L4* correlates with low expression of genes assigned to skin development and differentiation (**Fig. 1D**). These data indicate *MAB21L4* expression is reduced in cSCC tissue and induced during epidermal differentiation, suggesting a potential new role for *MAB21L4* in these processes.

**Fig. 1.**
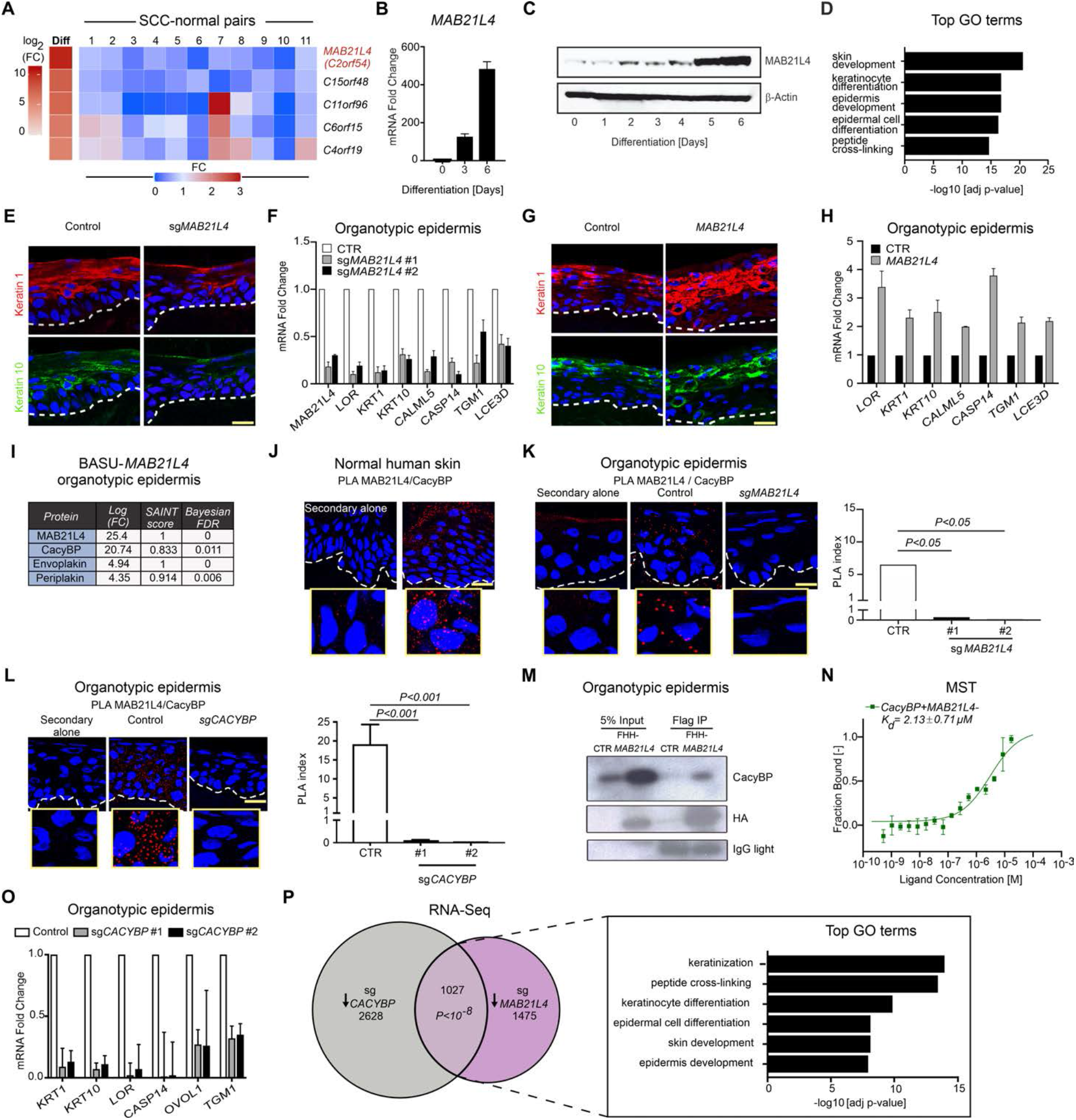
MAB21L4 is a regulator of epidermal differentiation that binds CacyBP. (**A**) RNA-seq heatmap of uncharacterized ORFs upregulated in primary human keratinocytes after six days of calcium-induced differentiation compared to undifferentiated, sub-confluent keratinocytes as control. The expression fold change (FC) of each ORF in 11 primary human cSCC with individual-matched normal skin as control is also shown. (**B**) MAB21L4 transcript induction during keratinocyte differentiation by qRT-PCR; undifferentiated cells (day 0) and days 3 and 6 of calcium-induced differentiation. (**C**) MAB21L4 protein induction over six days of calcium-induced differentiation in primary keratinocytes. (**D**) GO terms of genes with low expression when *MAB21L4* expression is also low using data from 27,247 publicly available microarray datasets. *P* values were adjusted using the Benjamini-Hochberg procedure with an FDR of 0.05. (**E**) Representative expression of differentiation proteins in *MAB21L4*-ablated (sg*MAB21L4*) human skin organoids generated by CRISPR/Cas9-mediated genetic disruption compared to a non-targeting control; differentiation proteins detected by immunofluorescence in red (keratin 1) and green (keratin 10), dotted line indicates basement membrane zone. (**F**) Differentiation gene mRNA quantitation in human skin organoids generated using two independent *MAB21L4* sgRNAs compared to control sgRNA. (**G**) Representative expression of differentiation proteins in human skin organoids generated from keratinocytes transduced to express MAB21L4; differentiation proteins detected by immunofluorescence in red (keratin 1) and green (keratin 10), dotted line indicates basement membrane zone. (**H**) Differentiation gene mRNA quantitation in human skin organoids with enforced expression of *MAB21L4* compared to a control vector. (**I**) Top MAB21L4-proximal proteins identified by BioID performed in human skin organoids. (**J**) PLA of endogenous MAB21L4 and CacyBP in normal human adult skin. PLA was performed in the absence of primary antibodies as a control (Secondary alone). (**K**) PLA of endogenous MAB21L4 and CacyBP in human skin organoids generated from *MAB21L4*-ablated primary keratinocytes (sg*MAB21L4*) compared to control sgRNA. Quantitation of PLA index with two independent *MAB21L4* sgRNAs was calculated for three fields of view using the number of dots per nucleus. (**L**) Representative PLA of endogenous MAB21L4 and CacyBP in human skin organoids generated from *CACYBP*-ablated primary keratinocytes (sg*CACYBP*) compared to control sgRNA and quantitation with two independent *CACYBP* sgRNAs. (**M**) Western blot analysis of FLAG co-immunoprecipitation of empty vector (CTR) or FLAG-HA-6xHIS (FHH)-MAB21L4 with endogenous CacyBP in human epidermal organoids. Data from three pooled technical replicates are shown. (**N**) Dissociation curve of recombinant purified CacyBP and MAB21L4 proteins measured by MST. By comparison, no binding was detected between CacyBP and MAFB, a protein of similar size to MAB21L4. Data show technical triplicates and are representative of results obtained in two independent experiments. (**O**) Differentiation gene mRNA quantitation in human skin organoids generated using two independent *CACYBP* sgRNAs compared to control sgRNA. (**P**) Venn diagram illustrating overlap between downregulated (↓) genes in *CACYBP*-ablated (n=3 biological replicates) and *MAB21L4*-ablated organotypic epidermis (n=2 biological replicates). Top GO terms associated with the 1,027 genes downregulated when *CACYBP* or *MAB21L4* are deleted in organotypic epidermis. *P* values were adjusted using the Benjamini-Hochberg procedure. All human skin organoids were grown for three days on dermis. Data in **F**, **H**, **K**, **L**, and **O** are shown as mean ± s.d. and are representative of three independent experiments unless otherwise specified. Statistical significance in **K** and **L** was determined using a two-tailed t-test. All scale bars, 100 μm.

To define *MAB21L4* function in the epidermis, we investigated the effects of MAB21L4 loss and gain in three-dimensional human skin organoids, a setting that faithfully recapitulates the architecture and gene expression profile of normal skin (17). CRISPR/Cas9-mediated genetic disruption with two independent single guide RNAs (sgRNAs) was used to create pools of *MAB21L4*-ablated primary human keratinocytes that were placed onto human dermis and grown for three days at the air-liquid interface to generate organotypic skin tissue. Deletion of *MAB21L4* attenuated the expression of both early (*KRT1*, *KRT10*, *CALML5*, *CASP14*) and late (*TGM1*, *LOR*, *LCE3D*) differentiation markers in regenerated skin tissue without affecting epidermal stratification (**Fig. 1E-F; Supplementary Fig. S1B**). Conversely, enforced expression of *MAB21L4* resulted in increased expression of these genes (**Fig. 1G-H; Supplementary Fig. S1C**). These results together illustrate that MAB21L4 is required for epidermal differentiation.

To gain insight into the molecular mechanisms that underlie MAB21L4’s impacts on differentiation, we used proximity-dependent biotin identification (BioID) to nominate MAB21L4’s protein interaction partners in human skin organoids. *MAB21L4* was fused to BASU, a Bacillus subtilis-derived biotin ligase (18). Primary human keratinocytes transduced to express the BASU-MAB21L4 fusion protein at levels approximating endogenous MAB21L4 expression or BASU only as control were used to generate human skin organoids. Following biotin labeling, streptavidin pull-down, and mass spectrometry analysis, the top MAB21L4-proximal proteins based on fold change over control and significance analysis of interactome (SAINT) score (19) included those known to be essential for epidermal barrier function, such as envoplakin and periplakin (20–22), as well as proteins that are uncharacterized in skin, such as calcyclin-binding protein (CacyBP), also known as Siah-1 interacting protein (SIP) (**Fig. 1I**). Proximity ligation assays (PLA) confirmed MAB21L4 co-localization with envoplakin (**Supplementary Fig. S1D**) and periplakin (**Supplementary Fig. S1E**).

We next focused on CacyBP, which is known to interact with members of the S100 family, cytoskeletal proteins, and components of E3 ubiquitin ligases (23), but whose expression and activity have not been studied in skin. PLA was used to confirm the physical proximity of endogenous MAB21L4 and CacyBP in adult human skin (**Fig. 1J**). Genetic ablation of either *MAB21L4* or *CACYBP* in human skin organoids abolished MAB21L4-CacyBP fluorescent PLA signal (**Fig 1K-L**). We then generated human skin organoids from keratinocytes transduced to express epitope-tagged MAB21L4 and demonstrated its association with CacyBP by PLA (**Supplementary Fig. 1F**) as well as co-immunoprecipitation (**Fig. 1M**). Microscale thermopheresis (MST) was used to define the CacyBP and MAB21L4 interaction in a quantitative manner with a Kd of 2.1 ± 0.7 µM and no binding was detected between CacyBP and MAFB, a negative control protein with a molecular weight similar to MAB21L4 (**Fig. 1N**). These data establish the physical proximity of MAB21L4 and CacyBP in three-dimensionally intact human skin tissue.

Given the ability of MAB21L4 to regulate epidermal differentiation, we next investigated whether CacyBP might play a similar role in this process. We confirmed a previously unrecognized requirement for CacyBP in epidermal differentiation by demonstrating impaired expression of differentiation markers in *CACYBP*-ablated human skin organoids that recapitulated the effects of MAB21L4 depletion (**Fig. 1O**; **Supplementary Figs. 1G-H**). To more fully investigate the relationship between MAB21L4 and CacyBP, we then analyzed *MAB21L4*- or *CACYBP*-deficient human skin organoids by RNA sequencing. This effort distilled a set of 1,027 coordinately downregulated genes enriched with those assigned to keratinization, keratinocyte differentiation, and skin development gene ontology (GO) terms (**Fig. 1P**). Enrichment of peptide cross-linking genes that encode transglutaminases and multiple cornified envelope components essential for the mechanical integrity and barrier function of the skin was also observed. These results identify a direct binding relationship between MAB21L4 and CacyBP and demonstrate that both proteins are essential for epidermal differentiation.

### MAB21L4 and CacyBP suppress invasion in cSCC organoids

To examine MAB21L4’s role in cancer, we first confirmed reduced MAB21L4 expression in cSCC as well as actinic keratoses (AK), the earliest clinically detectable cSCC precursors (**Fig. 2A; Supplementary Fig. S2A**). These data are consistent with the downregulation of *MAB21L4* observed at the mRNA level in cSCC and suggest that loss of MAB21L4 expression occurs early in cSCC development. We next investigated whether *MAB21L4* is subject to somatic copy number alterations (SCNAs) as genes frequently inactivated through deletion may function as tumor suppressors. SCNAs were analyzed in 9,927 pairs of tumor and matching normal genomes in 27 human cancer types in The Cancer Genome Atlas (TCGA) data set after filtering out genomic segments containing known tumor suppressor genes. In multiple TCGA cancer types, including cervical and lung SCC, *MAB21L4* was present at the nadir of the peak formed by aligning regions of somatic copy number deletions and also placed in a focal peak of deletion by the GISTIC algorithm (**Fig. 2B**) (24). Analysis of MAB21L4 expression in cSCC as well as lung (LUSC) and head and neck (HNSC) SCC, two TCGA epithelial malignancies for which mRNA expression of large numbers of both tumor and normal tissues is available, showed decreased expression of MAB21L4 in tumors and provided additional support for somatic *MAB21L4* loss in cancer (**Supplementary Fig. S2B**). We then performed Cox proportional hazard analysis of *MAB21L4* expression in a combined stratified epithelial cancer cohort composed of TCGA cervical (CESC) as well as HNSC and found that each unit of increased MAB21L4 expression was revealed to result in an additional 6% decrease in the hazard for death (**Supplementary Fig. S2C and Supplementary Table S1**). Taken together, these data support a model whereby MAB21L4 functions as a tumor suppressor in multiple epithelial tissues from distinct anatomical sites.

**Fig. 2.**
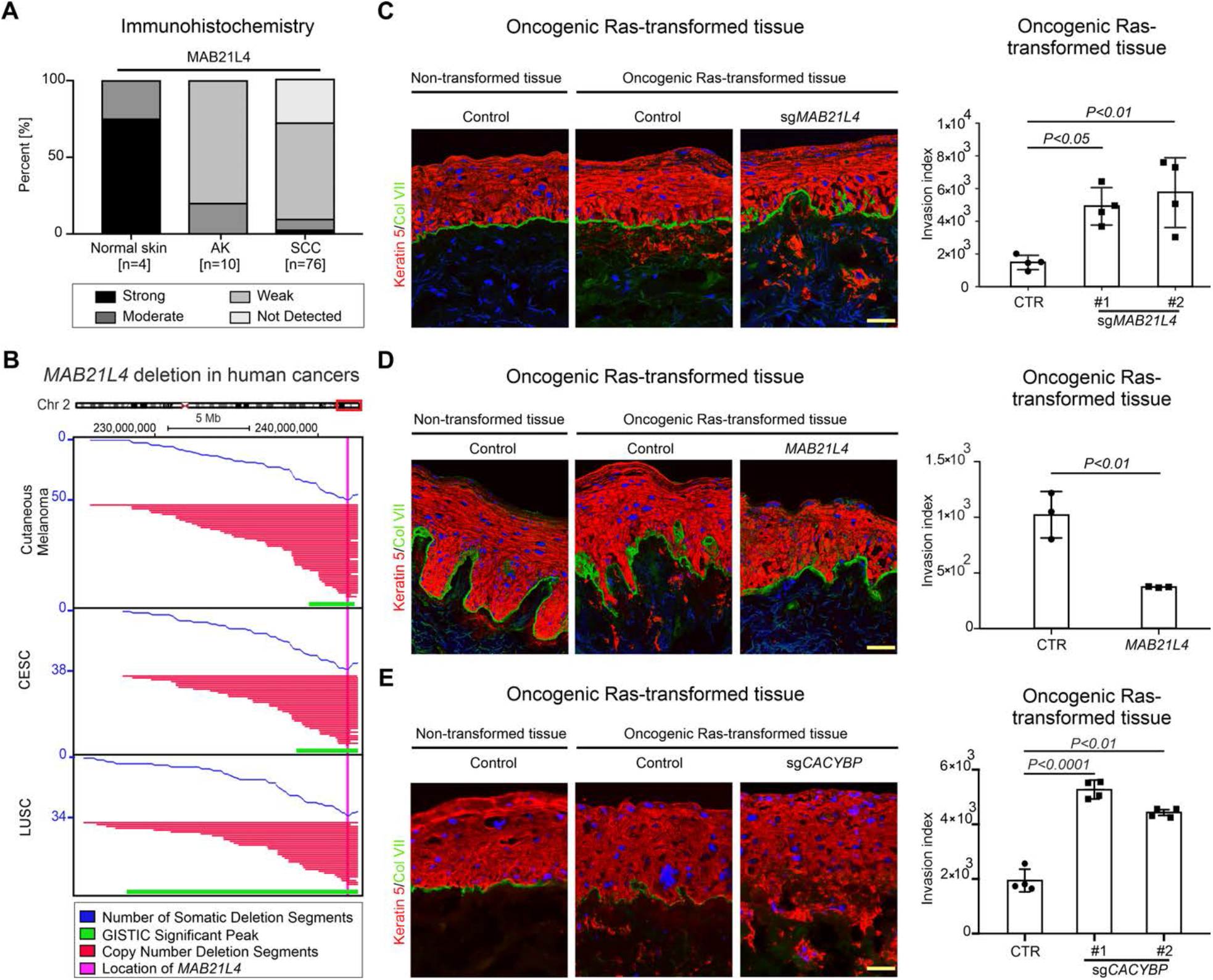
MAB21L4 and CacyBP suppress invasive human epidermal neoplasia. (**A**) MAB21L4 expression in normal skin, pre-cancerous actinic keratoses (AK), and cSCC. (**B**) Somatic deletion of *MAB21L4* in TCGA LUSC, CESC, and cutaneous melanoma (SKCM) datasets. Somatic deletion segments that do not span known tumor suppressor genes are shown in red. GISTIC-identified peak regions are shown in green. The position of the *MAB21L4* locus is shown in pink. (**C**) Normal primary human keratinocytes seeded onto dermis do not breach the epithelial boundary defined by the basement membrane (left); however, oncogenic Ras-expressing keratinocytes (keratin 5, red) invade into the stroma in this model (middle). Depletion of MAB21L4 using CRISPR/Cas9-mediated genetic disruption (sg*MAB21L4*) potentiates invasion (right). Quantitation of invasion index is shown with two independent *MAB21L4* sgRNAs, calculated as the integrated density of keratin 5 staining beneath the basement membrane divided by the length of the basement membrane in four fields of view. (**D**) Normal human skin organoids (left) are compared to their oncogenic Ras-transformed counterparts in the absence (middle) and presence (right) of enforced MAB21L4 expression. Quantitation of invasion index was calculated using three fields of view. (**E**) Loss of CACYBP by CRISPR/Cas9-mediated genetic disruption (sg*CACYBP*) enhances Ras-induced neoplastic invasion. Invasion index was quantitated with two independent *CACYBP* sgRNAs using four fields of view. Results were confirmed in at least two independent experiments with technical triplicates. All scale bars, 100 μm. Invasion index is shown as mean ± SD; significance was calculated using a two-tailed t-test.

To investigate MAB21L4 function in the cancer setting, we generated human skin organoids programmed by oncogenic H-Ras and cyclin dependent kinase Cdk4 to transform into invasive epidermal neoplasia that possesses the histology and global gene expression profile of human cSCC (25). *MAB21L4* expression is decreased in cSCC organoids compared to non-transformed skin tissue, consistent with its downregulation in human cSCC (**Supplementary Fig. S2D**). Genetic ablation of *MAB21L4* with two independent sgRNAs potentiated Ras-driven neoplastic invasion, as evidenced by an increase in the number of keratinocytes present beyond the epidermal-dermal boundary (**Fig. 2C**). Enforced expression of MAB21L4 had the reverse effect and suppressed neoplastic invasion as well as basement membrane degradation (**Fig. 2D and Supplementary Fig. S2E**). We then xenografted Ras-transformed human cSCC organoids that overexpressed MAB21L4 or a control vector onto immune-deficient mice and confirmed that MAB21L4 effectively blocks neoplastic progression in vivo (**Supplementary Fig. S2F**). Furthermore, *CACYBP* ablation with two independent sgRNAs enhanced neoplastic invasion in human cSCC organoids compared to a non-targeting sequence, recapitulating the effects of MAB21L4 depletion (**Fig. 2E**). These loss- and gain-of-function studies in cSCC organoids demonstrate a tumor-suppressive role for MAB21L4 in the skin that hinges upon MAB21L4-CacyBP interaction.

### MAB21L4/CacyBP-dependent regulation of RET expression

We next explored the mechanism of MAB21L4’s tumor-suppressive action by examining its dependent genes and compared the differentially expressed genes identified in human skin organoids with *MAB21L4* overexpression, *MAB21L4* or *CACYBP* gene ablation, and human cSCC. Genes that were downregulated by enforced *MAB21L4* expression and upregulated upon its deletion were selected to search for cancer-promoting genes that might be therapeutically actionable. The resulting gene set was then overlapped with the list of genes upregulated by *CACYBP* loss as well as those with increased expression in our series of 11 cSCC. This analysis revealed that the intersection of these four datasets contained the proto-oncogene *RET*, an RTK with established cancer relevance (**Supplementary Fig. S3A**). We confirmed differential expression of *RET* in these conditions, including an independent series (n = 73) of human cSCC (**Supplementary Fig. S3B-D**). Further support for RET activation in cSCC was provided by enrichment of a combined RET/oncogenic MAPK transcriptional signature (26) in tumors compared to normal skin (**Supplementary Fig. S3E**).

We then wished to determine which transcription factors (TFs) might be responsible for the observed changes in *RET* expression. GeneHancer was used to distill a list of TFs with binding sites within the RET promoter (27) and a TF enrichment analysis was subsequently performed with ChEA3 (28) focusing on these candidates and using upregulated genes in human cSCC and either *MAB21L4-* or *CACYBP-*ablated human skin organoids as input (**Supplementary Fig. S3F**). Examination of ENCODE chromatin immunoprecipitation (ChIP) sequencing (ChIP-seq) data (29) for the top TFs predicted to regulate *RET* transcription identified a distinct binding peak for ZFP69B, GLIS1, and CTCF in the *RET* promoter region. Notably, overexpression of *ZFP69B* and *GLIS1* has been reported in human cancer types known to possess RET-altered subsets (30, 31). These data together nominate candidate TFs for RET upregulation.

RTK signaling is also known to be tightly controlled by ubiquitylation, and previous studies have indicated that casitas B-lineage lymphoma (CBL) family as well as neural precursor expressed, developmentally downregulated 4 (NEDD4) E3 ligases promote RET ubiquitylation and degradation, although this process has not been characterized in skin (32–34). CacyBP in particular is known to interact with Siah1, the critical component of a multiprotein E3 ubiquitin ligase complex that targets β-catenin for destruction upon p53 induction (35), prompting us to investigate whether RET expression in the skin might be similarly regulated. After confirming the physical proximity of endogenous MAB21L4 and RET (**Supplementary Fig. S3G**), we used PLA to demonstrate that deleting *MAB21L4* with two independent sgRNAs reduced the fluorescent signal between CacyBP and RET in human skin organoids (**Fig. 3A**).

**Fig. 3.**
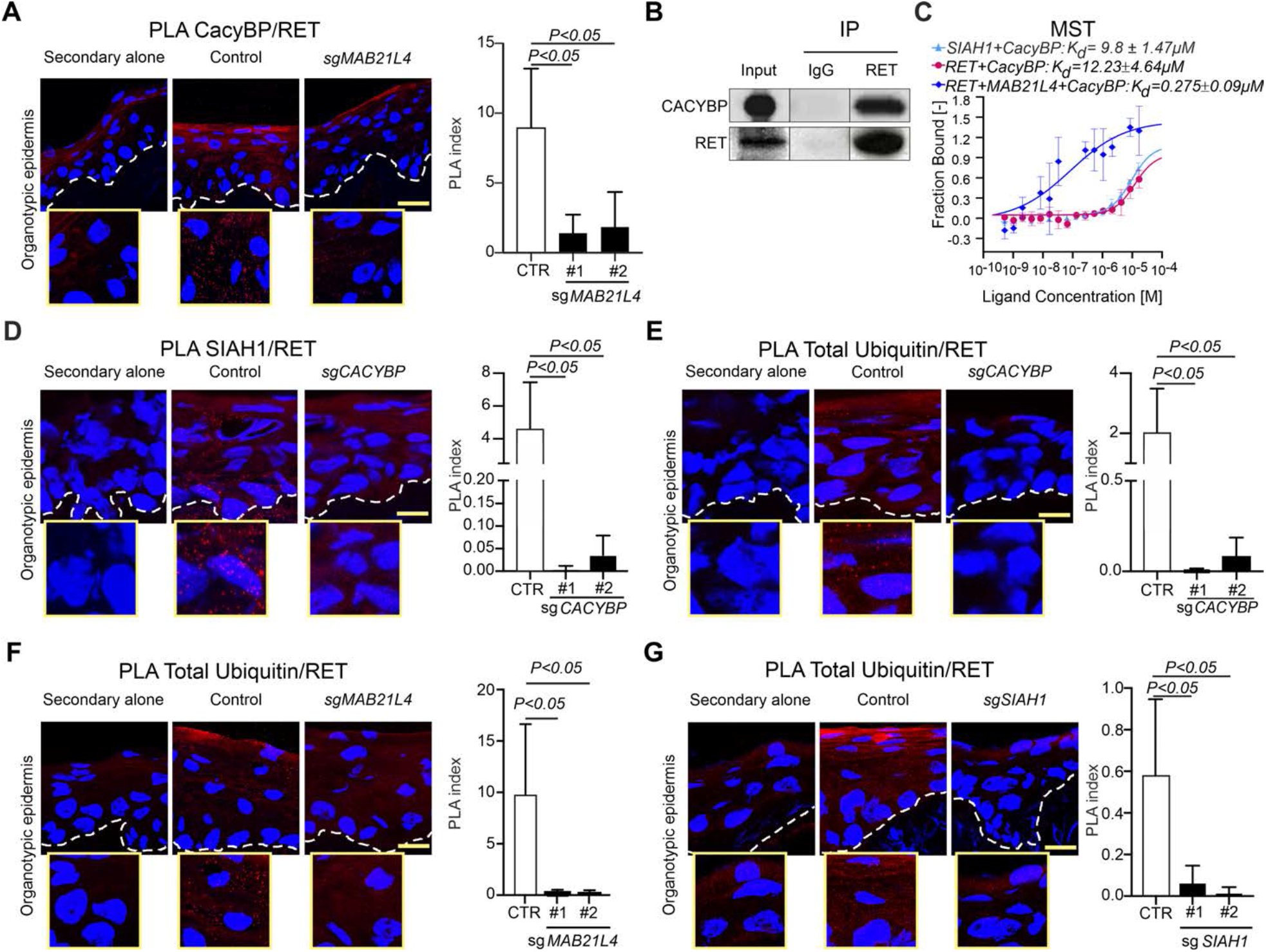
MAB21L4-CacyBP interaction controls RET proximity and ubiquitination by Siah1. (**A**) PLA of endogenous CacyBP and RET in human skin organoids generated from *MAB21L4*-ablated primary keratinocytes (sg*MAB21L4*) compared to control sgRNA. Quantitation of PLA index with two independent *MAB21L4* sgRNAs. (**B**) Western blot analysis of SCCIC1 cells subjected to RET or IgG co-immunoprecipitation. Data shown is representative of n=3 technical replicates. (**C**) Dissociation curve of recombinant purified CacyBP and RET proteins in the presence or absence of MAB21L4 measured by MST. Data show technical triplicates and are representative of results obtained in two independent experiments. PLA of endogenous Siah1 and RET in *CACYBP*-ablated human skin organoids (**D**); endogenous ubiquitin and RET in *CACYBP*-ablated (**E**), *MAB21L4*-ablated (**F**), and *SIAH1*-ablated (**G**) human skin organoids. In each case, quantitation of PLA index with two independent sgRNAs is shown. All organoids were grown for three days on dermis and results are representative of at least two independent experiments. All scale bars, 100 μm. PLA index was calculated from three fields of view using the number of dots per nucleus. Data represent mean ± s.d.; statistical significance was determined by two-tailed t-test.

Additional support for CacyBP interaction with RET was provided by co-immunoprecipitation and MST, with the latter demonstrating a 44-fold increase in binding affinity between CacyBP and RET when MAB21L4 was also present (**Fig. 3B-C**). Siah1 was run in parallel as a positive control for MST and bound to CacyBP with a Kd of 9.8 ± 1.5 µM, consistent with previously published values for this interaction (36). These data together support a direct interaction between CacyBP and RET that requires MAB21L4 and suggest a model wherein MAB21L4 functions as an adaptor protein that enforces proximity between CacyBP and RET.

We then used PLA to demonstrate the proximity of CacyBP-interacting Siah1 to RET in human skin organoids. The fluorescent PLA signal generated by Siah1 and RET proximity was markedly attenuated in *CACYBP*-ablated epidermal tissue and a similar reduction in RET-ubiquitin proximity was also observed (**Fig. 3D-E**). Loss of either *MAB21L4* or *SIAH1* also abolished the PLA signal generated by RET and ubiquitin proximity (**Fig 3F-G; Supplementary Fig. S3H**). Conversely, RET immunoprecipitated from human cells that overexpress MAB21L4 exhibited increased ubiquitination compared to cells transfected with a control plasmid (**Supplementary Fig. S3I**). Enforced expression of MAB21L4 also resulted in more rapid degradation of RET following the addition of cycloheximide to block protein synthesis (**Supplementary Fig. S3J**). Thus, ubiquitination of RET by the multiprotein Siah1-CacyBP E3 ligase complex is destabilizing and requires MAB21L4.

### Genetic ablation or inhibition of RET promotes differentiation and attenuates cancer progression in cSCC organoids

Given MAB21L4’s loss in multiple cancer types and its role in RET ubiquitylation, we reasoned that aberrant RET activation might drive cSCC as well as other related tumors. Increased expression of RET was confirmed in a second cohort of cSCC (37) as well as TCGA HNSC and elevated levels of *DUSP6* and *SPRY4*, two MAPK target genes that serve as biomarkers of RET activity (38), were also observed in both malignancies (**Fig. 4A and Supplementary Fig. S4A**). High expression of *RET* in a combined stratified epithelial TCGA cancer cohort (CESC and HNSC) predicted worse overall survival, with each unit increase in *RET* mRNA expression resulting in an 11% increase in hazard (**Supplementary Fig. S4B and Supplementary Table S1**). We then analyzed whole exome data from 53 cSCC-normal pairs (6, 39) and examined copy number calls in the TCGA HNSC cohort. *RET* copy number gains were detected in 36.8% (19 of 53) cSCC and 10.9% (57 of 522) HNSC, indicating that gene amplifications may contribute to *RET* overexpression. Somatic point mutations in *RET* were also identified in 14% (14 of 100) of cSCC (6) and 2.0% (10 of 522) HNSC, suggesting an alternate mechanism of *RET* activation in a subset of these malignancies consistent with that seen in medullary thyroid carcinomas and the hereditary cancer syndrome multiple endocrine neoplasia type 2 (MEN 2) (40). Recurrence of mutations was not observed; however, cSCC-associated *RET* variants included R844L and S409Y, which have been implicated in cancer (41). Further support for the pathogenicity of these mutations was provided by the Combined Annotation Dependent Depletion (CADD) framework (42), which assigned median C scores of 23.6 and 23.2 to cSCC-and HNSC-associated *RET* variants respectively, corresponding to the top 0.43% and 0.47% of deleterious mutations in the human genome and comparable to the scores of known oncogenic variants (**Fig. 4B and Supplementary Fig. S4C**). These data together demonstrate aberrant *RET* activation in cSCC as well as other stratified epithelial cancers.

**Fig. 4.**
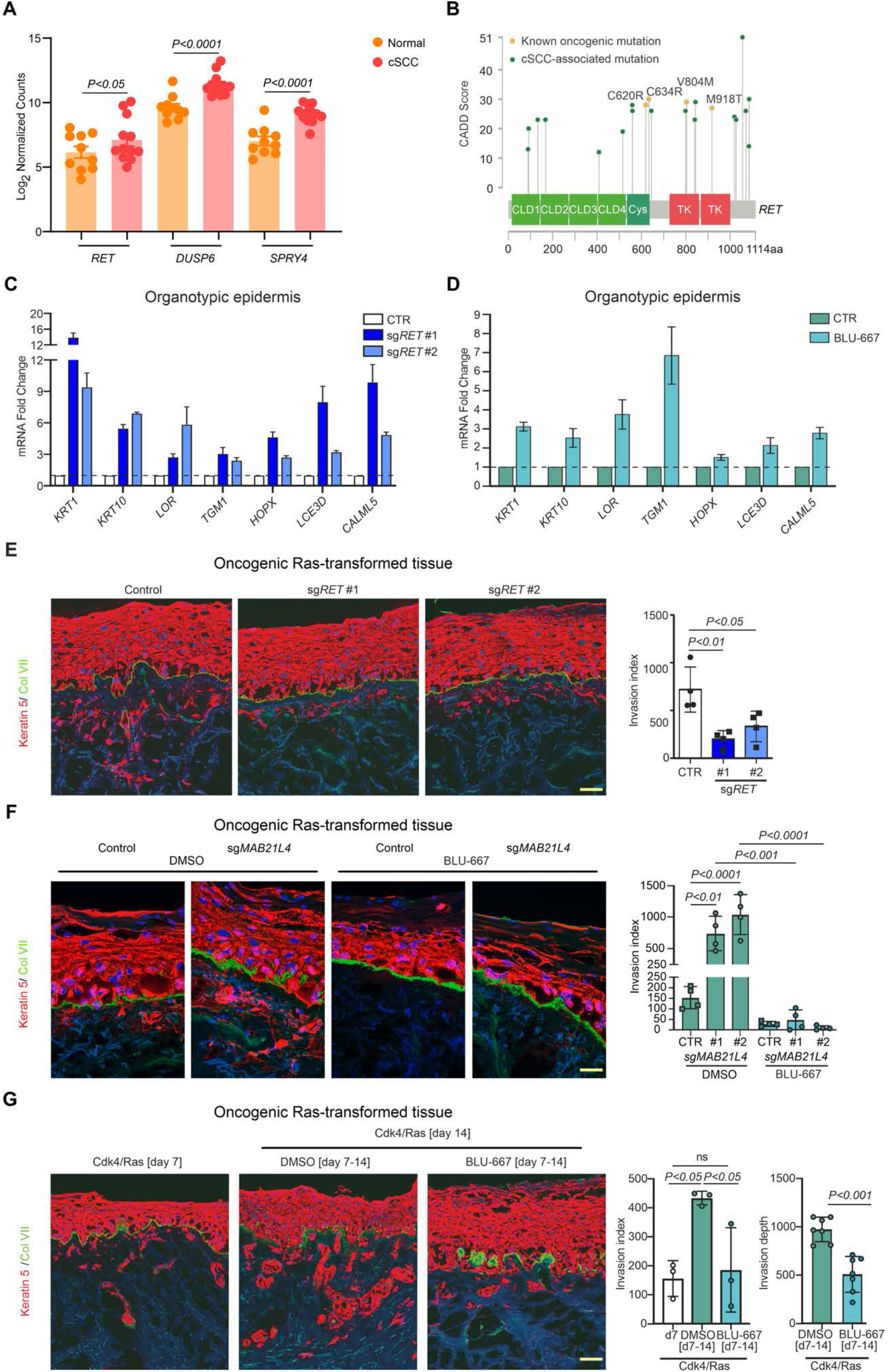
RET activation in cSCC promotes tumorigenesis in organoids and maintains keratinocytes in the undifferentiated state. (**A**) Increased mRNA expression of *RET* as well as *DUSP6* and *SPRY4*, two biomarkers of RET activity, in cSCC compared to normal skin. *P*-values were calculated using DESeq2. (**B**) cSCC-associated *RET* mutations are plotted as a function of CADD score and compared to known oncogenic mutations. CLD, cadherin-like domain. Cys, cysteine-rich domain. TK, tyrosine kinase. (**C**) Differentiation gene mRNA induction in human skin organoids generated using two independent *RET* sgRNAs compared to control sgRNA and after three days of growth on dermis. (**D**) Relative expression of differentiation genes in human skin organoids grown for three days on dermis in the presence or absence of BLU-667 (15 nM). (**E**) Keratin 5 (red) and collagen VII (green) immunofluorescence of oncogenic Ras-transformed human skin organoids generated from *RET*-ablated human primary keratinocytes (sg*RET*) or control sgRNA after 7 days of growth on dermis. Quantitation of invasion index using four fields of view from each of two independent *RET* sgRNAs. (**F**) Keratin 5 (red) and collagen VII (green) immunofluorescence of oncogenic Ras-transformed human skin organoids generated from *MAB21L4*-ablated human primary keratinocytes (sg*MAB21L4*) or control sgRNA after 7 days of growth on dermis; tissue was regenerated in the presence or absence of BLU-667, a highly selective RET inhibitor. Quantitation of invasion index in using four fields of view from each of two independent *MAB21L4* sgRNAs. (**G**) Keratin 5 (red) and collagen VII (green) immunofluorescence of oncogenic Ras-transformed human skin organoids after 7 or 14 days of growth on dermis; BLU-667 or DMSO was initiated on day 7. Quantitation of invasion index and invasion depth using at least three fields of view. Data in **E**, **F**, and **G** are shown as mean ± s.d. and are representative of at least two independent experiments. The invasion index is calculated from the integrated density of keratin 5 staining beneath the basement membrane divided by the length of the basement membrane. The invasion depth is measured as the depth of the tumor front below the basement membrane Bars in **C**-**D** represent the mean of three technical replicates ± s.d. and similar results were confirmed in two independent experiments. Statistical significance was determined by two-tailed t-test. All scale bars, 100 μm.

RET is most strongly expressed in the basal layer of the epidermis, suggesting a role in progenitor cell maintenance (**Supplementary Fig. S5A**). To test this directly, we disrupted *RET* using two independent sgRNAs to create pools of gene-targeted primary keratinocytes and placed these cells onto dermis to generate human skin organoids (**Supplementary Fig. S5B**). *RET* ablation had the opposite effect of *MAB21L4* or *CacyBP* deletion and resulted in increased expression of differentiation markers without affecting epidermal stratification (**Fig. 4C and Supplementary Fig. S5C**). Human skin organoids treated with the RET inhibitor BLU-667 similarly exhibited enhanced differentiation gene expression (**Fig. 4D and Supplementary Fig. S5D**). These results indicate a functional requirement for RET in maintaining the undifferentiated state characteristic of progenitor keratinocytes and are consistent with data in Drosophila demonstrating Ret expression in adult intestinal epithelial progenitor cells, but not their differentiated counterparts (43).

Loss of differentiation is often coupled to other cancer-enabling behaviors and BLU-667 is currently approved to treat *RET*-driven lung and thyroid cancers (44), prompting us to consider whether aberrant *RET* activation might be a targetable dependency in cSCC. We first demonstrated that *RET* ablation with two independent sgRNAs reduced cancer cell invasion in human cSCC organoids compared to a non-targeting sequence (**Fig. 4E**). We then generated cSCC organoids using *MAB21L4*-intact or *MAB21L4*-ablated primary human keratinocytes and maintained these neoplastic tissues in culture medium containing BLU-667 to further investigate the contribution of RET to cSCC development. Treatment with BLU-667 prevented basement membrane degradation and suppressed invasion accelerated by *MAB21L4* loss (**Fig. 4F**). BLU-667 was also able to prevent further invasion of keratinocytes when added to cSCC organoids after early tumorigenesis occurred, with fewer keratinocytes observed below the basement membrane and a reduction in the depth of tumor cell infiltration into the dermis (**Fig. 4G**). Further, topical administration of BLU-667 to skin organoids undergoing malignant transformation similarly reduced cancer cell invasion (**Supplementary Fig. S5E**). These data together demonstrate the effectiveness of targeting RET in cSCC development and progression.

### Anti-tumor effects of RET inhibition in cSCC and HNSC in vitro and in vivo

To further explore the effects of RET inhibition, we next evaluated the invasiveness of cSCC cells in the presence or absence of BLU-667. A431 and SCCIC1 cell invasion through Matrigel was suppressed by BLU-667 (**Supplementary Fig. S5F-G**). Multicellular tumor spheroids were then generated from these and two HNSC cell lines, SCC4 and SCC47, to assess their sensitivity to BLU-667. After confirming suppression of RET signaling by BLU-667 in the selected cell lines (**Figs. 5A-D; Supplementary Fig. S6A-D**), tumor spheroids were created by seeding cells into ultra-low attachment plates for 24 hours before transitioning to culture medium containing BLU-667 for 48 hours. RET inhibition by BLU-667 reduced cell viability in all tumor spheroids tested and disrupted aggregation at the outer edge, as evidenced by loss of compaction (**Fig. 5E-L**). BLU-667 treatment also induced differentiation gene expression in these cell lines, further confirming RET is required to maintain cells in the undifferentiated state (**Supplementary Fig. S6E-F**). These results provide orthogonal experimental support of RET dependency in cSCC and suggest the same therapeutic vulnerability is present in HNSC.

**Fig. 5.**
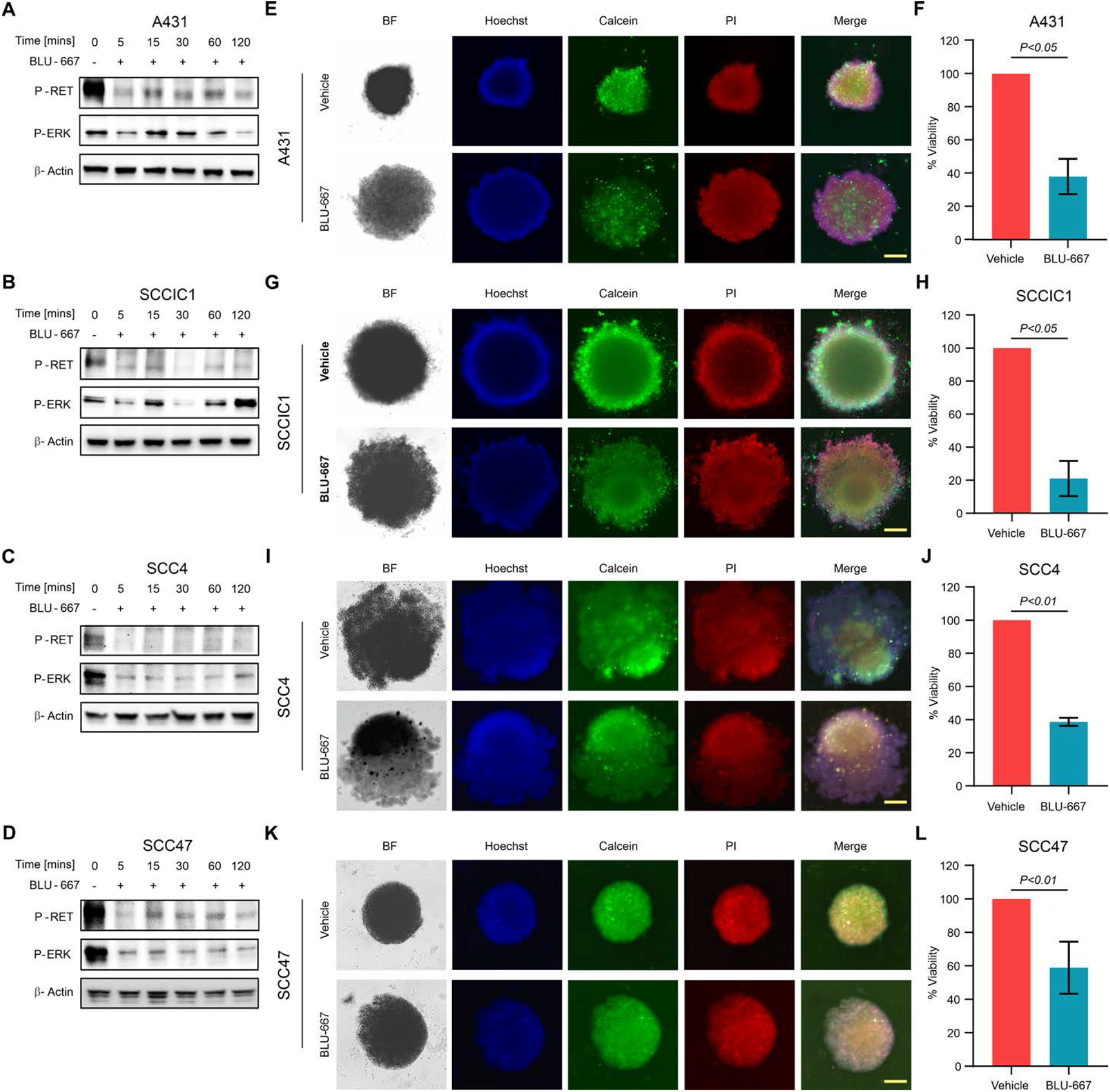
RET inhibition reduces SCC spheroid viability. Western blot analysis of A431 (**A**), SCCIC1 (**B**), SCC4 (**D**), and SCC47 (**D**) cells treated with BLU-667 showing decreased phospho-RET and phospho-ERK. (**E**, **G**, **I**, **K**) Bright-field (BF) and immunofluorescence images of SCC spheroids. PI, propidium iodide. All scale bars, 100 μm. (**F**, **H**, **J**, **L**) Quantitation of cell viability in SCC spheroids. Data represent the mean of eight technical replicates ± s.d. and similar results were confirmed in three independent experiments.

We next assessed the effectiveness of selective RET inhibition on cSCC tumorigenesis in vivo. Human A431 subcutaneous xenografts were established in NCG mice and daily treatment with BLU-667 was initiated after tumor engraftment. While the size of vehicle-treated cSCC xenografts continued to increase, tumor growth plateaued upon BLU-667 treatment (**Fig. 6A-C**). A reduction in RET signaling was observed in BLU-667 tumor lysates compared to vehicle controls, suggesting RET pathway inhibition underlies slowed cSCC growth (**Supplementary Fig. S7A**). Fewer Ki67-positive tumor cells were present in BLU-667-treated cSCC and provided additional confirmation of decreased cell proliferation in vivo (**Fig. 6D**). Upon further histologic examination, we noted BLU-667 treatment was associated with the presence of numerous keratin debris-filled cystic structures characteristic of differentiating keratinocytes (**Fig. 6E**). Compared to vehicle-treated cSCC, BLU-667-treated tumors also exhibited higher expression of differentiation markers consistent with a transition from a proliferative progenitor state to a post-mitotic one (**Fig. 6F and Supplemental Fig. S7B**). These results parallel our SCC organoid and in vitro studies demonstrating disrupted cancer progression and concomitant differentiation by BLU-667.

**Fig. 6.**
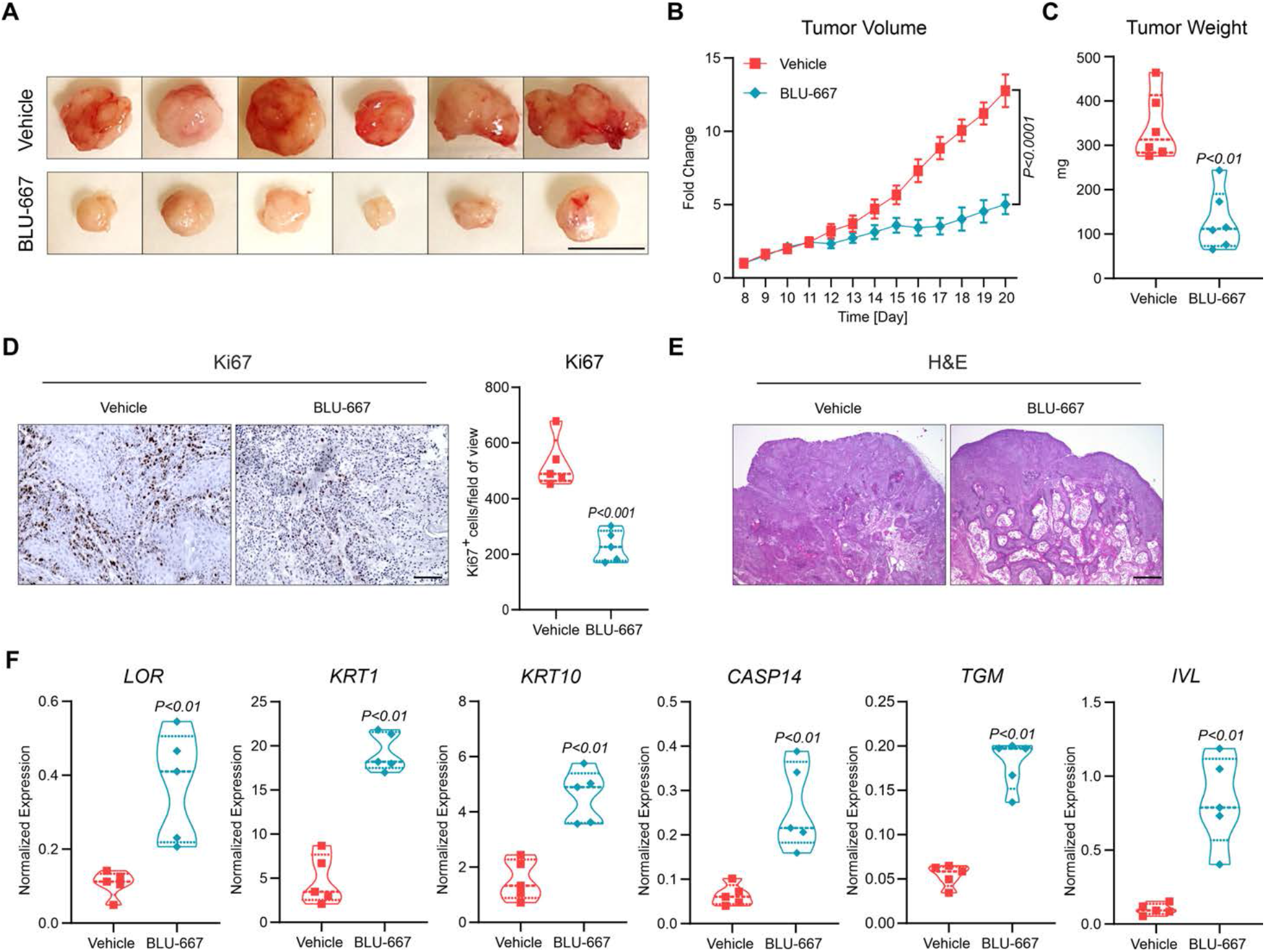
Inhibition of RET suppresses tumorigenesis and induces differentiation in vivo. (**A**) A431 xenograft tumors. (**B**) Antitumor activity of BLU-667 compared to vehicle showing the response of A431 xenografts (n=6 in each treatment group) as a fold change in relative tumor volume. Data represent the mean ± s.e.m. *P* value was determined using a two-way ANOVA. (**C**) Antitumor activity of BLU-667 compared to vehicle showing the response of A431 xenografts (n=6 in each treatment group) as tumor weight measured on the last day of treatment. Data represent the mean ± s.d. *P* value was determined using a two-tailed t-test. (**D**) Representative Ki67 immunohistochemistry of A31 xenografts treated with BLU-667 compared to vehicle. Quantitation of Ki67-positive cells from 5 fields of view. Dashed lines indicate the median and the first and third quartiles are represented by the dotted lines. *P* value was determined using a two-tailed t-test. (**E**) Representative hematoxylin and eosin staining of A431 xenografts treated with BLU-667 compared to vehicle. (**F**) qRT-PCR analysis of differentiation gene expression in A431 xenografts treated with BLU-667 or vehicle (n=5 in each treatment group). Dashed lines indicate the median and the first and third quartiles are represented by the dotted lines. *P* value was determined using a two-tailed t-test. Scale bar is 1 cm in **A** and 100 μm in all other panels.

## Discussion

Few non-invasive medical therapies are available to the increasing number of individuals with cSCC who decline or are not candidates for surgical treatment. Our study demonstrates the therapeutic potential of targeting the RET receptor tyrosine kinase in cSCC using human skin organoids, tumor spheres, and a xenograft mouse model. Increased RET expression in cSCC as well as other stratified epithelial cancers including HNSC is associated with poor survival, and 14% of cSCC as well as a subset of HNSC harbor mutations predicted to alter protein function. Functionally, we demonstrate that genetic disruption of *RET* or selective RET kinase inhibition both suppress tumorigenesis while concomitantly enhancing differentiation, suggesting a model wherein RET is required to maintain the proliferative undifferentiated keratinocyte cell state that becomes predominant in neoplasia (**Fig. 7**).

**Fig. 7.**
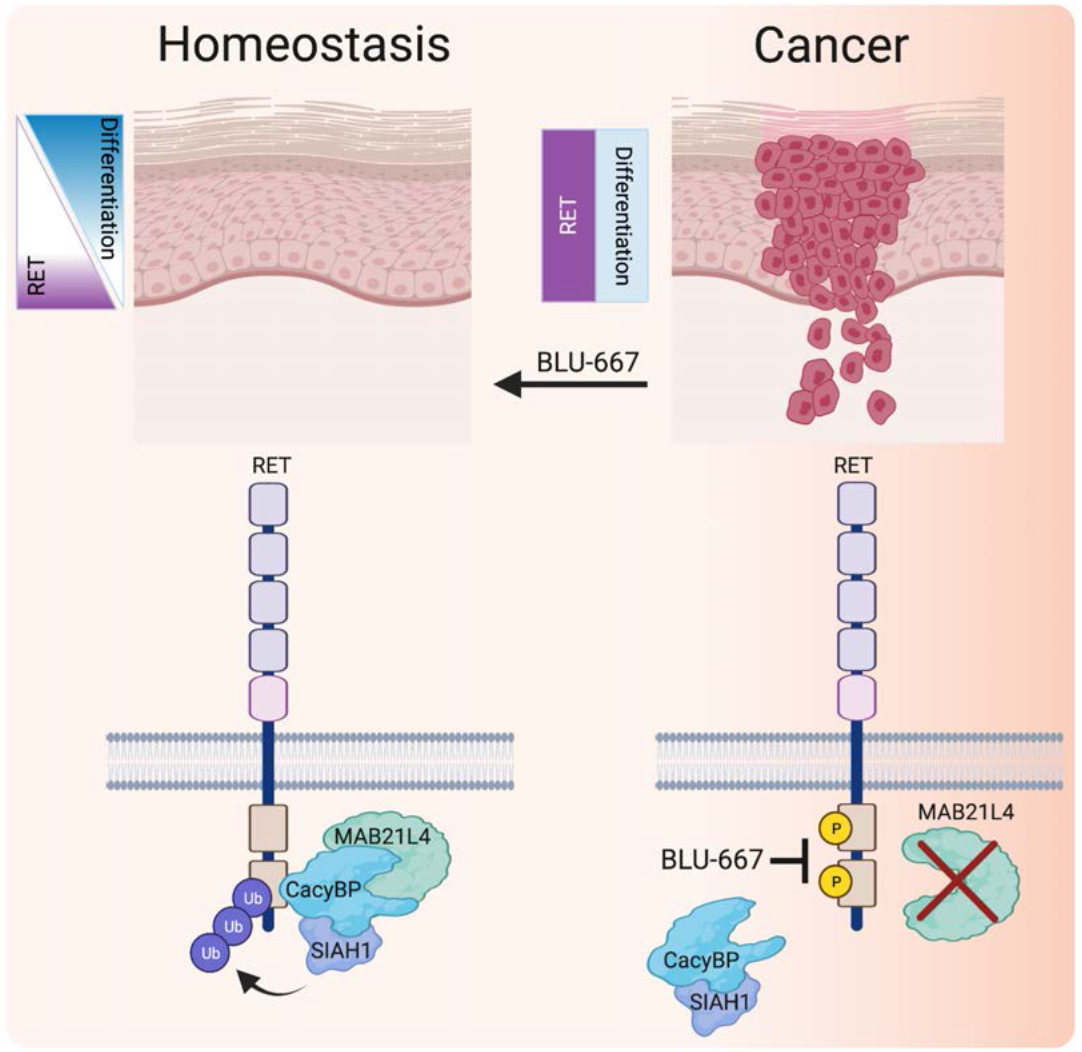
MAB21L4-CacyBP regulation of RET expression. Proposed model: Loss of MAB21L4-CacyBP interaction in cSCC enables sustained RET activation and disrupts the balance between proliferation and differentiation.

RET activation in thyroid and non-small cell lung cancer is commonly achieved through activating point mutations as well as gene rearrangements (45) and may also result from overexpression of the wild type protein, such as occurs in breast cancer (46). Our findings in cSCC and HNSC suggest these malignancies are also RET-driven and are consistent with the mechanisms of RET activation established in other cancers. A functional role for RET in cSCC development has not previously been described, although *RET* fusions have been reported at low frequencies in spitzoid melanocytic neoplasms (47) and *ret* transgenic mice are born with systemic melanosis from which spontaneous melanomas capable of metastasis arise (48).

The function of RET is vital to early development, as demonstrated by knockout studies in mice that reveal impacts on kidney organogenesis and gut innervation as well as germ cell survival in the developing male testis (49, 50). *RET* inactivation has not been studied directly in human skin and our results indicating this favors differentiation are consistent with previous studies in Drosophila (43). Our findings are also in agreement with reports demonstrating RET activation engages integrin signaling (51), which is known to negatively regulate epidermal differentiation (52). Taken together, these data suggest pharmacologic RET inhibition suppresses cSCC tumorigenesis by re-establishing the normal keratinocyte differentiation program and may explain the effectiveness of β1 integrin blockade in attenuating human epidermal neoplasia mimicking SCC (53).

BLU-667 is a once-daily selective oral RET inhibitor that targets wild type RET as well as activating RET mutations and fusions. It is currently approved to treat RET-positive metastatic non-small cell lung cancer or advanced thyroid cancer and appears to have a more favorable safety profile compared to previously approved multi-kinase inhibitors due to its high selectivity for RET over other kinases, such as VEGFR (54).

Though generally well-tolerated, 40% of patients who received BLU-667 in the phase I/II ARROW trial required dose reductions and 4-6% discontinued treatment due to adverse events that included neutropenia and anemia (54). Topical administration would reduce the risks of systemic treatment while enabling higher local levels of inhibition at the tumor site, and we confirmed the feasibility of this approach by applying BLU-667 topically to evolving human cSCC organoids. Further studies are needed to verify and fully characterize the safety and clinical benefit of topical BLU-667 as well as other RET inhibitors in cSCC tumorigenesis.

Our data indicate that the level of RET expression is controlled by MAB21L4, a protein of previously unknown function that enforces differentiation and suppresses tumorigenesis. *MAB21L4* is named for its resemblance to the *mab-21* gene, which acts cell autonomously to control tail pattern in C. elegans males as well as non-autonomously to specify cell fate outside of the tail region (55, 56). Three homologs of *mab-21*, termed mab-21-like 1 (*MAB21L1*), mab-21-like 2 (*MAB21L2*), and mab-21-like 3 (*MAB21L3*) have been characterized in vertebrates that together control early embryogenesis and differentiation, impacting eye, head, neural tube, and body wall development (57–59). With the exception of *mab21l3*, which has been shown to control differentiation in Xenopus embryonic epidermis (59) and is only expressed at low levels in human epidermis (60), mab-21 proteins have not previously been functionally characterized in the skin. Interplay between other mab-21 proteins and RET may exist and additional studies are warranted to define their precise impacts on RET signaling. Furthermore, the presence of *MAB21L4* somatic deletions or its decreased expression in multiple cancer types, including CESC and HNSC where loss of MAB21L4 predicts worse overall survival, suggests a broader role for MAB21L4 that extends beyond the skin.

Our results show that a physical proximity relationship between MAB21L4 and CacyBP underlies RET-mediated keratinocyte progenitor maintenance and tumor-like behaviors such as migration and cancer cell invasion. CacyBP is a critical component of a ubiquitin E3 ligase complex known to ubiquitylate β-catenin through interaction with Siah1 (35) and appears to control RET stability in a similar manner. Interestingly, transcriptional upregulation of *RET* was also detected in our analysis comparing *MAB21L4*- and *CACYBP*-dependent genes to those upregulated in human cSCC. The TFs responsible for *RET* upregulation and their contribution to RET expression merit further investigation.

In summary, our study establishes RET activation as an enabler of SCC carcinogenesis targetable by pharmacologic inhibition and presents a rationale for further investigation into the clinical use of selective RET inhibitors in this setting. High RET expression is orchestrated by loss of MAB21L4, an early event occurring in precancerous AK that disrupts RET proximity to CacyBP-Siah1, thereby preventing its ubiquitylation. Early post-approval data of BLU-667 in non-skin malignancies confirms its overall safety and tolerability; as such, we anticipate that these and related compounds, particularly those that can be delivered to the skin topically, may be a valuable addition to existing treatment options for cSCC and its precursor lesions.

## Online Methods

### Accession Numbers

The data discussed in this publication have been deposited in the NIH Gene Expression Omnibus and are accessible through accession number GSE190907.

### Human Tissue Samples

All human tissues were studied under an IRB-approved protocol and all patients provided written informed consent. All samples were validated by histological analysis.

### Cell Culture

All cells were maintained at 37°C in 5% CO_2_. Primary human keratinocytes and fibroblasts were isolated from freshly discarded skin surgical specimens as previously described (25). Keratinocytes were cultured in a 1:1 mixture of Medium 154 (Life Technologies M-154-500) supplemented with human keratinocyte growth supplement (HKGS) containing 0.2% bovine pituitary extract (BPE), 5 µg/ml bovine insulin, 0.18 µg/ml hydrocortisone, 5 µg/ml bovine transferrin, and 0.2 ng/ml human epidermal growth factor (EGF) with Keratinocyte-SFM (Life Technologies 17005-142) supplemented with recombinant human EGF and BPE. Keratinocyte differentiation was induced in vitro by growing the cells at full confluence with 1.2 mM calcium added to the culture medium for up to 6 days. Fibroblasts, HEK-293T, Phoenix, and A431 cells were maintained in DMEM medium (Gibco) supplemented with 10% FBS. SCCIC1 cells were cultured in DMEM/F-12 50/50 medium (Corning 10-092-CV) with L-glutamine and 15 mM HEPES containing 10% serum and growth factors as previously described (61). SCC4 and SCC47 cells were grown in DMEM/F-12 50/50 (Corning 10-092-CV) medium supplemented with 10% FBS. All lines tested negative for Mycoplasma contamination and cancer cell lines were confirmed to be authentic and unique by short tandem repeat fingerprinting.

### Human Skin Organoids

Generation of human skin organoids was performed as previously described (17). Briefly, 1×10^6^ keratinocytes were seeded onto a 1 cm^2^ section of devitalized human dermis and cultured at the air-liquid interface for up to 6 days for differentiation studies. To generate organotypic invasive epidermal neoplasia, devitalized human dermis was populated with human primary fibroblasts at least 1 week prior to keratinocyte seeding. Keratinocytes were transduced with oncogenic HRas as well as Cdk4 and seeded onto fibroblast-containing dermis. Neoplastic tissues were harvested for analysis up to 14 days post-seeding.

### Lentiviral Transduction

Lenti-X 293T cells were transfected with FLAG-HA-6xHIS (FHH)-tagged *MAB21L4* or an empty FHH vector along with the helper plasmids pCMV-dR8.91 and pUC-MDG to generate lentiviral medium. Human primary keratinocytes were transduced with FHH-*MAB21L4* or FHH only with 5 µg/ml polybrene. Two days post-transduction, keratinocytes were trypsinized and plated in a larger culture flask in growth medium supplemented with 1 µg/ml puromycin. Cells were selected in puromycin for 2 days or until mock-transduced control cells were completely eradicated.

### Retroviral Transduction

Phoenix cells were transfected with plasmids containing the cDNA sequences of *CDK4* or *HRAS* to generate retroviral medium (53). Human primary keratinocytes were transduced with retroviral medium supplemented with 5 µg/ml polybrene by centrifugation at 1000 rpm for 1 hour at room temperature. The retroviral medium was replaced with fresh growth medium immediately after centrifugation and cells were returned to 37°C in 5% CO2.

### CRISPR/Cas9 Gene Ablation

Potential sgRNA sequences predicted to have high sensitivity as well as specificity were generated using a CRISPR design tool (62) and cloned into the pLentiGuide plasmid (63). Concentrated lentiviral medium was generated for each sgRNA as well as Cas9 using the pLEX_Cas9 plasmid (63) as described above. For gene ablation studies, human primary keratinocytes from two different donors were pooled and transduced with Cas9 lentiviral medium. Two days post-transduction, the cells were trypsinized and plated in a larger culture flask in growth medium supplemented with 2 µg/ml blasticidin. Cells were selected in blasticidin for 2 days and then re-seeded and transduced with sgRNA lentiviral medium. Two days post-sgRNA transduction, the cells underwent selection in growth medium supplemented with 1 µg/ml puromycin. For the *CACYBP*-ablation studies, keratinocytes immortalized by retroviral transduction with the pLXSN16 plasmid encoding the HPV E6 and E7 proteins served as the starting epithelial cell pool. The following sgRNA sequences were used: Control (GACCGGAACGATCTCGCGTA); *MAB21L4* sg1 (GCAGCACGTTCTCTGCGCGC); *MAB21L4* sg2 (GAGTAGTCCACGATGAAGCGG); *CACYBP* sg1 (TTATCTGACTGATCCCATCC); *CACYBP* sg2 (ACTTACCTCTTCTGAAGCCA); *RET* sg1 (TCCTCGTCGTACACGGTCAC); *RET* sg2 (TGGCGTACTCCACGATGAGG); *SIAH1* sg1 (TCATAGGCGACGATTGACTT); *SIAH1* sg2 (TTCCCGGCGCCGAGACCGAC).

### Immunofluorescence

Human skin organoids were embedded in Optimal Cutting Temperature (OCT) compound (Sakura Finetek) and 7 µm tissue sections were mounted onto polylysine slides (Thermo Scientific). Tissue cryosections were fixed in 100% methanol at −20°C for 10 min and subsequently permeabilized by incubation with 0.1% Triton X-100 (Sigma-Aldrich) in PBS. The tissues were then incubated for 30 minutes at room temperature with blocking buffer (horse serum diluted 1:10 in PBS) and then for 1 hour with primary antibodies diluted in 0.05% Triton X-100 (Sigma-Aldrich) and 1:100 horse serum in PBS. The primary antibodies used were: MAB21L4 (1:50, Biorbyt, orb159238), Collagen Type VII (1:50, Millipore, MAB2500), Keratin 1 (1:500, BioLegend, 905201), Keratin 10 (1:500, Thermo Fisher Scientific, MS611P), CacyBP (1:50, Proteintech, 11745-1-AP), Keratin 5 (1:500, BioLegend, 905501). After 2 washes, the tissues were incubated for 45 minutes with secondary antibodies diluted in 0.05% Triton X-100 (Sigma-Aldrich) and 1:100 horse serum in PBS. The secondary antibodies used were: goat anti-mouse IgG Alexa Fluor 488 (1: 500, Thermo Fisher Scientific, A-11001) and goat anti-rabbit IgG Alexa Fluor 594 (1: 500, Thermo Fisher Scientific, A-11012). The tissues were then mounted with Prolong Gold Antifade Mountant with DAPI (Thermo Fisher Scientific) and imaged using an LSM 880 Airyscan Fast Live Cell inverted confocal microscope (Zeiss). Images of several z-stacks were acquired and the maximum intensity projection image was generated by using the Fiji-ImageJ package (64).

### Proximity Proteomics in Human Skin Organoids

A triple hemagluttinin (3xHA) tag and BASU biotin ligase (18) were fused to the MAB21L4 protein in the pLEX vector. Primary human keratinocytes were transduced with 3xHA-BASU-MAB21L4 or 3xHA-BASU lentiviral medium as described above. Following selection in growth medium supplemented with 1 µg/ml puromycin, 1×10^6^ keratinocytes were seeded onto a 1 cm^2^ section of devitalized human dermis and cultured at the air-liquid interface for 5 days. On day 6 post-seeding, proximal proteins were labeled by adding 50 µM biotin to the growth medium for 16 hours. The epidermis from each tissue was separated from the dermis, minced with a scalpel, rinsed in PBS to remove biotin-containing growth medium, lysed in high-SDS RIPA buffer (Pierce), and then sonicated using a Bioruptor (Diagenode). The cell lysates were then centrifuged through Microsep columns (Pall) to deplete free biotin and then incubated with Dynabeads MyOne Streptavidin C1 beads (Thermo Fisher) at 4°C under rotation for 16 hours. Affinity purification was performed as previously described (65) and proteins were analyzed by LC-MS/MS using a Proxeon EASY nanoLC system (Thermo Fisher Scientific) coupled to an Orbitrap Elite mass spectrometer (Thermo Fisher Scientific). All mass spectra were analyzed with MaxQuant software version 1.5.5.1. MS/MS spectra were searched against the Homo sapiens Uniprot protein sequence database (version January 2017) and GPM cRAP sequences (commonly known protein contaminants). Precursor mass tolerance was set to 20 ppm and 4.5 ppm for the first search where initial mass recalibration was completed and for the main search, respectively. Product ions were searched with a mass tolerance 0.5 Da. The maximum precursor ion charge state used for searching was 7. Carbamidomethylation of cysteines was searched as a fixed modification, while oxidation of methionines and acetylation of protein N-terminal were searched as variable modifications. Enzyme was set to trypsin in a specific mode and a maximum of two missed cleavages was allowed for searching. The target-decoy-based false discovery rate (FDR) filter for spectrum and protein identification was set to 1%. Mass spectrometry proteomics data was analyzed for interactor candidates with SAINTq, using a SAINT probability score >0.867. Proteins were required to include at least one uniquely mapping peptide. Proteins that appeared in fewer than 60 out of 80 BioID experiments in the CRAPome, a database of negative controls and background contaminants, was used to further prioritize proteins (66).

### Proximity Ligation Assay (PLA)

PLA was performed using Duolink PLA reagents (Sigma-Aldrich, DUO92008) according to the manufacturer’s instructions. Briefly, 7 µm tissue cryosections were fixed in methanol as described above, followed by blocking with the provided blocking buffer and then incubated with primary antibodies against the two proteins of interest. The primary antibodies used for PLA analyses were: MAB21L4 (1:50, Biorbyt, orb159238), CacyBP (1:50, Novus Biologicals, H00027101-B01P), Envoplakin (1:50, Novus Biologicals, NBP2-37940), HA (1:50, BioLegend, 901501), Periplakin (1:50, LSBio, LS-C482731), CacyBP (1:50, Proteintech, 11745-1-AP), RET (1:50, NSJ Bioreagents, F53523), Ubiquitin (1:100, Abcam, ab7780), Siah1 (1:50, Genetex, GTX113268). After 2 washes, the tissues were incubated with the PLA probes anti-mouse MINUS (Sigma-Aldrich, DUO92004) and anti-rabbit PLUS (Sigma-Aldrich, DUO92002), followed by the ligation and amplification reactions. Tissues were then mounted with Prolong Gold Antifade Mountant with DAPI (Thermo Fisher Scientific) and imaged using a LSM 880 Airyscan Fast Live Cell inverted confocal microscope (Zeiss). Images of several z-stacks were acquired and the maximum intensity projection image was generated using the Fiji-ImageJ package.

### Co-Immunoprecipitation

Protein lysates from human skin organoids were extracted as described in the Western Blotting section. FHH-MAB21L4 protein was immunoprecipitated by incubating 95% of the lysate with anti-Flag M2 Affinity gel (Sigma-Aldrich) at 4°C under rotation for 16 hours. The gel beads were washed 3 times in cold TBS containing protease inhibitors (Sigma-Aldrich) and then eluted in 2X Laemmli Buffer (Bio-Rad) followed by western blotting. RET was immunoprecipitated from 293T cells by incubating 95% of the lysate with Dynabeads Protein G (Invitrogen, 10003D) conjugated to anti-RET antibody (Cell Signaling Technology, 3223).

### Quantitative RT-PCR

RNA was extracted using the RNeasy Plus Mini Kit (Qiagen) according to the manufacturer’s instructions and reverse transcribed to cDNA using iScript reverse transcriptase per the manufacturer’s protocol (Bio-Rad). Quantitative PCR was performed on a Roche LightCycler 480 II using the SYBR green kit (Fisher Scientific). The cycling parameters were: 94°C for 10 seconds, 60°C for 10 seconds, 72°C for 10 seconds, for a total of 45 cycles. Gene expression was quantified relative to the housekeeping genes RPL32 or GAPDH using the 2–ΔΔCt method 72. Primer sequences are listed in Supplementary Table S2.

### Western Blotting

Cells and organoids were both lysed in RIPA buffer (Thermo Fisher Scientific) supplemented with phosphatase inhibitor and protease inhibitor cocktails (Sigma-Aldrich). For human skin organoids, the epidermis was separated from the dermis, immersed in RIPA buffer, and subjected to 4 cycles of disruption using the gentleMACS Dissociator (Miltenyi Biotec). The primary antibodies used were: MAB21L4 (1:500, Abcam, ab185350); β-actin (1:1000, Sigma-Aldrich, A1978), HA (1:1000, Cell Signaling Technology, 3724S), HA (1:1000, BioLegend, 901501), CacyBP (1:500, Proteintech, 11745-1-AP), GAPDH (1:1000, Millipore, MAB374), GAPDH (1:1000, Cell Signaling Technology, 2118), Cdk4 (1:1000, Santa Cruz Biotechnology, sc-749), H-Ras (1:500, Santa Cruz Biotechnology, sc-520), RET (1:750, Cell Signaling Technology, 3223), p-RET (phospho-Y1062) (1:750, Abcam, ab51103), ubiquitin (1:500, Abcam, ab7780), Siah1 (1:50, Genetex, GTX113268), p-p44/42 MAPK (Erk1/2, phospho-T202/Y204) (1:1000, Cell Signaling Technology, 4370), p-Akt (phospho-S473) (1:1000, Cell Signaling Technology, 4060). The secondary antibodies were used at 1:10000 and were: IRDye 680RD goat anti-mouse (LI-COR, 926-68070), IRDye 800CW goat anti-mouse (LI-COR, 926-32210), IRDye 680RD goat anti-rabbit (LI-COR, 926-68071) and IRDye 800CW goat anti-rabbit (LI-COR, 926-32211). Proteins were visualized on a LI-COR Odyssey CLx. For chemiluminescence visualization, the secondary antibodies were: Amersham ECL HRP-conjugated donkey anti-rabbit (GE Healthcare, NA934) and Amersham ECL HRP-conjugated sheep anti-mouse (GE Healthcare, NA931). SuperSignal West Dura or SuperSignal West Femto maximum sensitivity substrate (Thermo Fisher Scientific) was used for protein visualization.

### RNA-seq

Human tissues were minced with a scalpel and then subjected to homogenization in RLT Plus buffer (Qiagen) with β-mercaptoethanol added in Lysing Matrix D tubes (MP Biomedicals) using a FastPrep homogenizer (MP Biomedicals). RNA extraction was otherwise performed using the RNeasy Plus Mini Kit (Qiagen) according to the manufacturer’s instructions. RNA integrity was verified using the RNA 6000 Nano Kit (Agilent) in combination with the 2100 Bioanalyzer (Agilent). mRNA libraries (polyA) were prepared and sequenced on the HiSeq 2500 (Illumina) with a target of 25 million mapped reads per sample. FASTQ data were aligned to the transcriptome using STAR version 2.5.4b69 and count data were generated with RSEM version 1.3.170 against the human genome assembly GRCh38. Differential gene expression analysis was performed using DESeq2 with a false-discovery rate (FDR) threshold of 0.171.

### Microscale Thermopheresis

All recombinant proteins were isolated from 293T cells. 10 µg of pLEX_FHH- or FLAG-tagged protein-of-interest plasmids were transfected per 10 cm plate of ∼70% confluent 293T cells using Lipofectamine 3000 (Thermo Fisher). After 48 hours, cells were washed with ice-cold PBS and harvested with lysis buffer (50 mM Tris HCl pH 7.4, 150 mM NaCl, 1 mM EDTA, 1% Triton X-100, 1X EDTA-free Protease Inhibitor Cocktail (Sigma) in Ultrapure water). Cells were incubated in lysis buffer for 30 min on ice with frequent vortexing, sonicated three times for 10 seconds at 10% power, then centrifuged at max speed for 10 min at 4°C to remove debris. The supernatant was collected and the protein concentration quantified by Pierce BCA protocol (Thermo Fisher). To bind/isolate the tagged proteins-of-interest, a saturating volume of anti-FLAG M2 affinity gel (Millipore Sigma) (1 mL 50/50 M2 slurry per 10 mg total protein) was washed with TBS (50 mM Tris HCl pH 7.4, 150 mM NaCl) then incubated with the lysate, diluted to 2 mg/mL, for 2 hours at 4°C. The beads were then rotated with Wash Buffer (50 mM Tris HCl pH 7.4, 3 mM EDTA, 0.5% IPEGAL, 10% glycerol, 500 mM NaCl, 1X EDTA-free Protease Inhibitor Cocktail (Sigma), 0.1mM DTT; Protease Inhibitor Cocktail and DTT were both added immediately before using the buffer) for 10 min at 4°C and quickly rinsed again in Wash Buffer. To elute the protein from the beads, the beads were equilibrated in an Elution Buffer wash (PBS, 0.05% Tween) then incubated overnight at 4°C in Elution Buffer with 0.5 mg/mL 3XFLAG peptide at four times the bead volume. The following day, the beads were spun down and the eluted volume was concentrated using 3k or 10k molecular weight cutoff Amicon columns (Millipore). The protein concentration of the purified, concentrated tagged proteins was quantified by running the sample against a BSA dilution series on a 10% Bis-Tris gel and staining with InstantBlue Protein Stain (Expedeon). To conduct the MST experiments, FHH-tagged target proteins (MAB21L4, RET, and Siah1) were labeled using RED-tris-NTA 2^nd^ generation dye from the Monolith His-Tag labeling kit (Monolith). Following the manufacturer’s protocol, FHH-tagged target protein was incubated in the dye at a 2:1 protein to dye ratio; once bound, the labeled target protein was added at a final concentration of 50 nM to a 16-point 1:1 PBST dilution series of the ligand protein (n-terminus FLAG-tagged CacyBP), with the highest starting concentration at 17.1 µM. FHH-tagged RET was also incubated with MAB21L4 for 15 minutes at room temperature prior to labeling and adding to the CacyBP dilution series. All 16 reaction mixtures were loaded into Monolith NT.115 Capillaries (NanoTemper Technologies), and MST was measured using a Monolith NT.115 instrument (NanoTemper Technologies) at room temperature. All runs were conducted at 40% MST power and 60% LED power. Signal to noise ratio and MST data from triplicate measurements were analyzed using MO.Affinity Analysis software (NanoTemper Technologies).

### Immunohistochemistry

Tissue microarrays containing samples of normal human skin and cSCC were purchased from Biomax US. AKs were collected under a protocol approved by our Institutional Review Board. Antigen retrieval was performed in 0.01 M Na Citrate (pH 6) and the following primary antibodies were used: MAB21L4 (1:500, Biorbyt, orb159238), RET (1:200, NSJ Bioreagents, F53523), KRT1 (1:200, Biolegend, 905201), LOR (1:200, Biolegend, PRB-145P-100). The secondary antibodies used were: goat anti-mouse IgG (ImmPRESS HRP Reagent Kit, Vector, MP-7452), goat anti-rabbit IgG (ImmPRESS HRP Reagent Kit, Vector, MP-7401), followed by the use of the chromogen DAB (ImmPACT DAB Peroxidase Substrate Kit, Vector, SK-4105). Analysis of immunohistochemistry staining was non-blinded given the distinct histologies of normal skin, AK, and cSCC.

### Drug Treatments

15 nM BLU-667 (Carbosynth) was added to the growth medium of human skin organoids for 7 days post-seeding or for 7 days starting one week after seeding. 0.1% DMSO was used as a vehicle control. Cycloheximide (Sigma) was dissolved in ethanol and added to A431 cells for 30 minutes at a final concentration of 100 µg/ml.

### Cell Invasion and Migration Assays

Matrigel invasion assays were performed using BioCoat Matrigel Invasion chambers with Matrigel matrix (Corning 354480). Cells were first starved for 24 hours by culturing in medium depleted of growth factors and then seeded into the insert in starving medium containing either 15 nM BLU-667 or 0.1% DMSO. The corresponding complete growth medium was placed in the well to establish the growth factor gradient and the plate was incubated at 37°C in 5% CO_2_ for 24 hours. Invasion was analyzed 24 (A431) or 48 hours (SCCIC1) post-seeding. For SCCIC1 cells, the growth medium in both the insert and well was refreshed 24 hours post-seeding to re-establish the growth factor gradient. At the time of invasion analysis, the insert was washed with PBS, fixed with 4% PFA for 30 min, washed again in PBS, and then stained with 0.01% crystal violet for 4 hours. The insert was then rinsed in tap water, gently scrubbed with a cotton swab on the upper surface, and then allowed to dry prior to imaging. Invading cells were counted per 2.5X field using Fiji-ImageJ.

### Tumor Spheroids

20,000 cells were seeded per well in a 96-well ultra-low attachment round bottom plate (Costar CLS7007) in their usual growth medium. After 24 hours of growth at 5% CO_2_ at 37°C, sphere formation was confirmed by bright field microscopy and the medium was transitioned to that containing 4.5 nM BLU-667 or 0.1% DMSO. Cell viability was measured qualitatively using the Live-Dead cell viability kit (Millipore) or quantitatively with the CellTiter-Glo 3D cell viability assay (Promega) following the manufacturer’s protocol.

### Statistical Analysis

To visualize somatic deletions at the MAB21L4 locus, publicly available TCGA datasets were downloaded from UCSC Xena (https://tcga.xenahubs.net:443) post-removal of common germline copy number variation. BEDTools was used to interrogate the MAB21L4 interval using <-0.25 as the threshold for significant copy number loss (67). Segments containing genes categorized as tumor suppressors by the COSMIC Cancer Gene Census (version GRCh37) were removed from analysis (68). The segments were then visualized using the UCSC genome browser. Segments were also used to plot a MAB21L4 deletion frequency percentage measured in 23 TCGA cancer types. For this analysis, SCNAs that overlap with COSMIC Cancer Gene Census tumor suppressors were removed.

To determine the effect of *MAB21L4* or *RET* expression on survival, TCGA expression data was downloaded from UCSC Xena. The prognostic potential of *MAB21L4* or *RET* was investigated in a combined stratified epithelial cancer cohort comprised of cervical cancer (CESC) as well as head and neck squamous cell carcinoma (HNSC). For each gene, Cox Proportional Hazards modeling was used to analyze the relationship between *MAB21L4* or *RET* mRNA expression and overall survival. The R modeling package rms 5.1-3.1 was used to fit the statistical models.

Venn diagrams were generated via eulerr (https://cran.r-project.org/package=eulerr) in R. For the 4-way Venn diagram in Figure S3A, a 0.2 TPM minimum filter was imposed across the replicates of each dataset. A TPM fold change threshold of 1.5 was used to call differential expression. Significance of overlap between separate categories was determined with a permutation resampling approach using 100,000 repetitions.

*RET*, *DUSP6*, and *SPRY4* expression in cSCC and HNSC compared to normal tissues was performed using publicly available RNA-seq data (15,37,69). The Combined Annotation Dependent Depletion (CADD) framework (version GRCh38-v1.6) was used to predict SNV deleteriousness. Gene set enrichment analysis was performed using fgsea version 1.18 (70) using a combined RET (DOI: 10.3180/R-HSA-8853659.2) and oncogenic MAPK (DOI: 10.3180/R-HSA-6802957.5) signature. Gene ontology analysis was done using ToppFun from the ToppGene Suite (71). Copy number alterations were detected using CNVkit version 0.9.8 (72) and GISTIC2.0 (24). GDC copy number parameters were used for GISTIC2.0.

### Animal Studies

All animal studies were performed in compliance with policies approved by the Stanford University Administrative Panel of Laboratory Animal Care. For skin xenograft studies, human cSCC organoids were first generated as above. After 5 days of growth at the air-liquid interface, tissues were grafted onto the dorsal backs of 6-week old female hairless SCID mice (Charles River, SHO, strain code 474) and then wrapped with petroleum-impregnated gauze followed by application of Tegaderm and Coban dressings (3M). Xenografts were unwrapped 7 days after initial grafting. For subcutaneous xenograft studies, 1.5×10^6^ A431 cells were suspended in 50 µL PBS and 50 µL Matrigel (Corning). Cells were injected subcutaneously into the flanks of 9-week-old female NOD CRISPR Prkdc Il2r Gamma mice (Charles River, NCG, strain code 572) with a 27-gauge needle and palpable tumors formed over 8 days. Mice were then divided into two groups to ensure an even distribution of tumor volume and injected once daily peritumorally with either vehicle alone (5% DMSO + 10% Solutol + 85% of 20% HPBCD) or 48 mg/kg BLU-667 diluted in the vehicle formulation. Tumor volumes were measured once daily using calipers, and treatment groups were blinded to measuring personnel. Tumor volume was estimated using the formula 4/3π(r1r2r3), where r1 is the radius of length, r2 is the radius of width and r3 is the radius of height. Mice were treated daily for 12 days before harvesting the tumors. As tumor weight is an objective measurement, the investigator was not blinded to the experimental group during this assessment.

## Acknowledgments

We thank Zurab Siprashvili, Ryanne A. Brown, Kerri Rieger, Brittany Stinson, Dan E. Webster, Shiying Tao, Bernd Jandeleit, Ishani Das, John Sunwoo, Bryan Sun, and Markus Kretz for reagents and helpful discussions. Confocal images were acquired in the Cell Sciences Imaging Facility (CSIF) at Stanford. Mass spectrometry was performed by the Proteomics Core at the Sanford-Burnham-Prebys Medical Discovery Institute. Histology was performed by Pauline Chu. The Translational Applications Science Center (TASC) at Stanford assisted with protein extraction from tissues. RNA-seq was performed by the Yale Center for Genome Analysis. The results published here are in part based upon data generated by the TCGA Research Network: https://www.cancer.gov/tcga. This paper is dedicated to Tom and Wei-Li Lee.

