## Supplementary Data for "RET activation controlled by MAB21L4-CacyBP interaction drives squamous cell carcinoma"

### Supplementary Figures


| | |
|---|---|
| Supplementary Fig. S1. | Expression of MAB21L4 and its proximal proteins in human epidermis. |
| Supplementary Fig. S2. | MAB21L4 expression in human cancers and organoids. |
| Supplementary Fig. S3. | Regulation of RET expression by MAB21L4 and CacyBP |
| Supplementary Fig. S4. | RET activation in cancer. |
| Supplementary Fig. S5. | RET maintains epidermal progenitor cells in an undifferentiated state and its inhibition suppresses tumor-enabling behaviors in cSCC. |
| Supplementary Fig. S6. | RET inhibition induces differentiation in SCC spheroids. |
| Supplementary Fig. S7. | BLU-667 inhibits RET signaling and enhances differentiation in vivo. |


### Supplementary Tables


| | |
|---|---|
| Supplementary Table S1. | Univariable relationship between gene expression and overall survival in TCGA cervical (CESC) and head and neck cancer (HNSC). |
| Supplementary Table S2. | Primer sequences. |

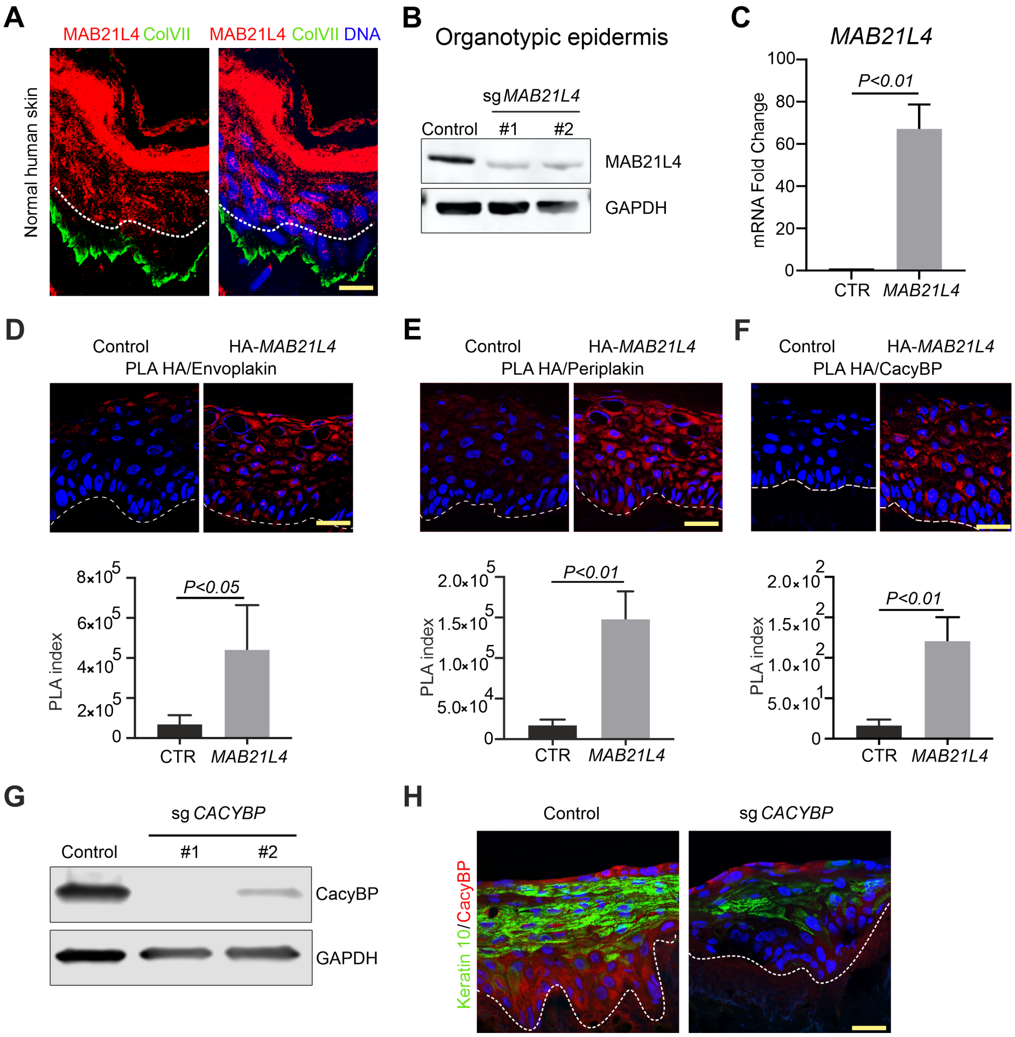


**Fig. S1. Expression of MAB21L4 and its proximal proteins in human epidermis.** (**A**) Expression of MAB21L4 in normal human adult skin. The dotted white line demarcates the basal layer from the suprabasal layers and collagen VII (green) decorates the basement membrane zone. (**B**) MAB21L4 expression in human skin organoids generated from *MAB21L4*-ablated primary keratinocytes via two independent sgRNAs (sg*MAB21L4*) compared to a non-targeting control sgRNA (Control). (**C**) *MAB21L4* mRNA quantitation in representative human skin organoids generated from keratinocytes transduced to express *MAB21L4*. Data represents n=3 technical replicates. PLA of HA and endogenous envoplakin (**D**), periplakin (**E**), and CacyBP (**F**) in human skin organoids generated from human primary keratinocytes transduced to express HA-MAB21L4 or an empty vector (Control) and grown for six days on dermis. PLA index in **D**, **E**, and **F** was calculated for three fields of view using the integrated density of signal per number of nuclei (envoplakin and periplakin) or the number of dots per nuclei (CacyBP). (**G**) Western blot analysis of CacyBP expression in epidermal organoids generated from *CACYBP*-ablated human primary keratinocytes via two independent sgRNAs (sg*CACYBP*) compared to a non-targeting control sgRNA (Control). (**H**) Expression of differentiation marker keratin 10 (green) in *CACYBP* (red)-ablated human skin organoids generated by CRISPR/Cas9-mediated genetic disruption compared to a non-targeting control after three days of growth on dermis; dotted line indicates basement membrane zone. Data shown are representative of three independent experiments. All scale bars, 100 μm. Bars in **C-F** represent the mean ± s.d. and statistical significance was determined using a two-tailed t-test.

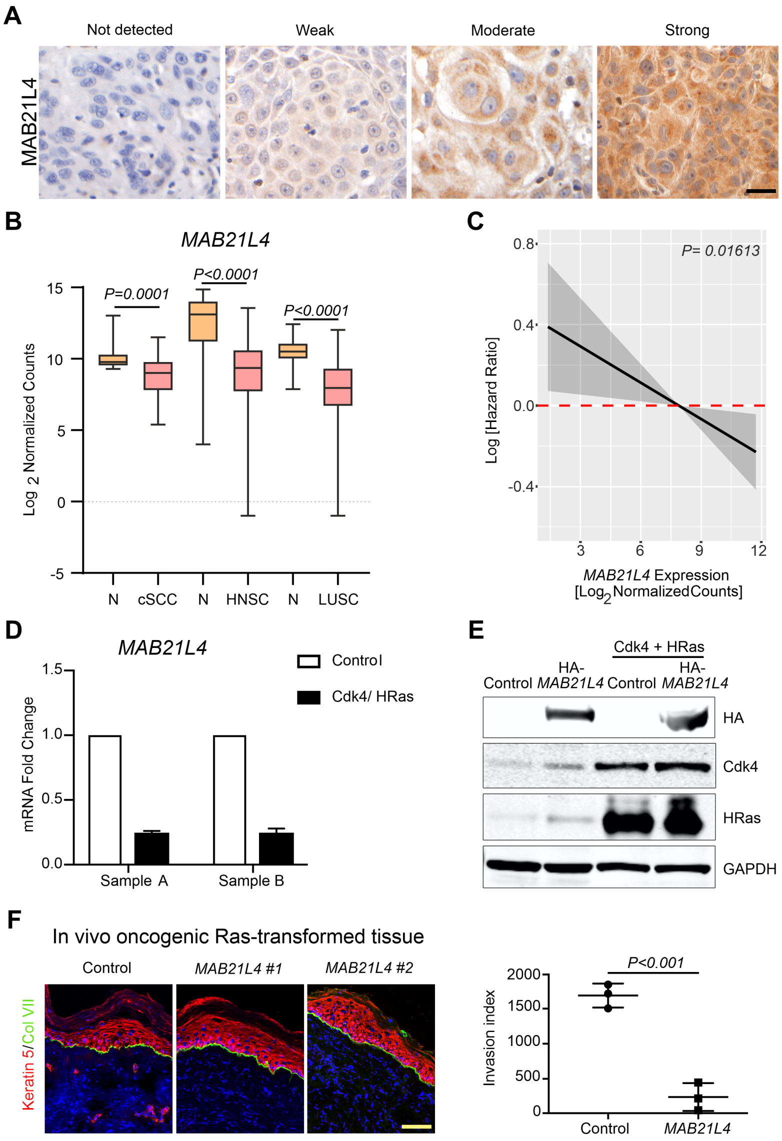


**Fig. S2. MAB21L4 expression in human cancers and organoids.** (**A**) Representative images of MAB21L4 immunohistochemical staining in cSCC tissues. (**B**) *MAB21L4* expression is decreased in cSCC as well as TCGA lung (LUSC) and head and neck (HNSC) SCC compared to normal tissues (N). Box shows twenty-fifth to seventy-fifth percentiles; whiskers show minimum to maximum, lines show medians. DESeq2 was used to compute *P*-values. (**C**) Cox hazard analysis of *MAB21L4* expression and survival in a combined stratified epithelial cancer cohort (TCGA CESC and HNSC). (**D**) *MAB21L4* transcript downregulation in human skin organoids programmed to transform into malignancy by oncogenic Cdk4/Ras and measured by qRT-PCR; sample A and B are biological replicates. Bars represent mean ± s.d., three technical replicates per experiment. (**E**) Western blot analysis of keratinocytes used to generate invasive human epidermal neoplasia. (**F**) Immunofluorescence of keratin 5 (red) and collagen VII (green) in xenografted epidermal tissues programmed to transform by oncogenic Cdk4/Ras; primary human keratinocytes were first transduced to express *MAB21L4* or an empty vector as control. Each experimental group contained n=3 mice and data show representative images from one Cdk4/Ras control xenograft as well as two different Cdk4/Ras/MAB21L4 xenografts. The invasion index was calculated using three fields of view (one field per mouse). Data are shown as mean ± s.d.; significance was calculated using a two-tailed t-test. All scale bars, 100 μm.

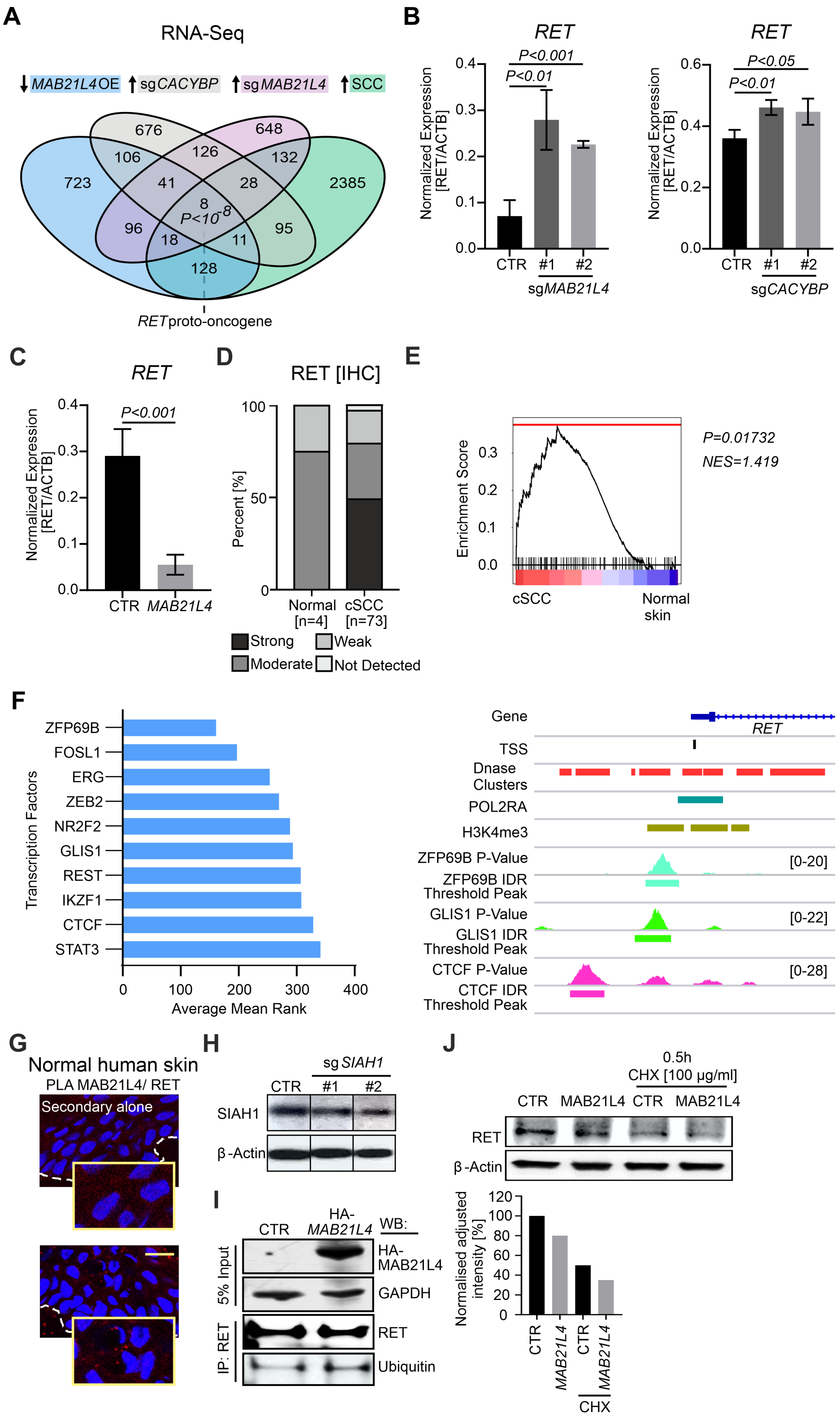


**Fig. S3. Regulation of RET expression by MAB21L4 and CacyBP.** (**A**) Venn diagram illustrating overlap between genes upregulated in 11 human cSCC compared to patientmatched normal skin as control (↑SCC), genes upregulated in *MAB21L4*-ablated human skin organoids (↑sg*MAB21L4*), genes downregulated when MAB21L4 is overexpressed in human skin organoids (↓MAB21L4 OE), and genes upregulated in *CACYBP*-ablated human skin organoids (↑sg*CACYBP*); the *P* value for the intersection of all four datasets is shown and was determined by bootstrap resampling. (**B**) *RET* transcript induction in *MAB21L4*- and *CACYBP*-ablated human skin organoids. Bars represent the mean ± s.d. of technical triplicates and significance was determined by two-tailed t-test. Similar results were obtained in at least two independent experiments. (**C**) *RET* transcript downregulation in human skin organoids with enforced *MAB21L4* expression. Bars represent the mean ± s.d. of technical triplicates and significance was determined by two-tailed t-test. (**D**) RET immunohistochemical staining in normal skin and cSCC. (**E**) Enrichment of the RET and oncogenic MAPK transcriptional signatures in RNA-seq expression data from cSCC compared to normal skin. NES, normalized enrichment score. Adjusted *P* value was calculated using the FGSEA package. (**F**) Potential regulators of *RET* were identified using ChEA3 to compare transcription factor enrichment in cSCC, *MAB21L4*-ablated human skin organoids, and *CACYBP*-ablated human skin organoids. IDR Threshold Peak and/or signal *P* value data from ENCODE ChIP-seq in HEK293 cells (ZFP69B, GLIS1, CTCF, and H3K4me3) or aggregated ENCODE cell types (DNase clusters and POL2RA) at the *RET* locus. (**G**) PLA of endogenous MAB21L4 and RET in normal human adult skin. PLA was performed in the absence of primary antibodies as a control (Secondary alone). Scale bar, 100 µm. (**H**) Western blot analysis of *SIAH1*-ablated keratinocytes. (**I**) Western blot analysis of ubiquitin expression in human HEK293T cells transduced with an empty vector (CTR) or HA-MAB21L4 and subjected to RET co-immunoprecipitation. (**J**) Western blot analysis and quantification using densitometry of RET expression in A431 cells transduced to express an empty vector (CTR) or MAB21L4 in the presence or absence of cycloheximide (CHX). Similar results were obtained in two independent experiments.

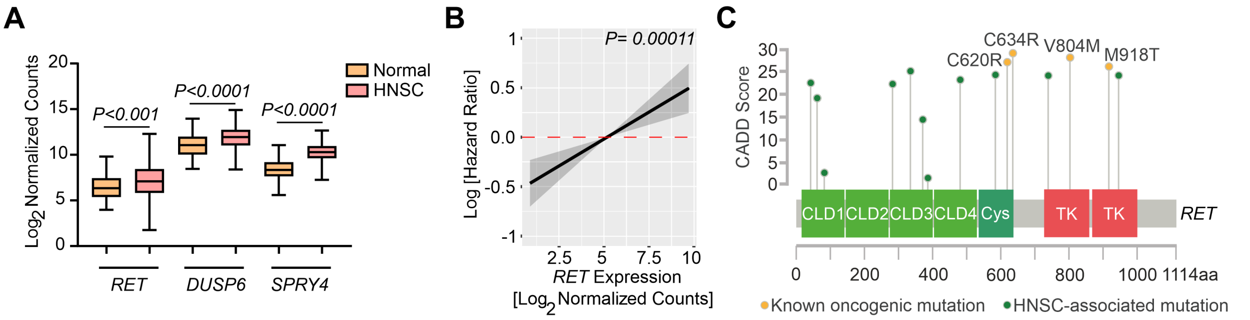


**Fig. S4. RET activation in cancer.** (**A**) *RET* expression is increased in TCGA HNSC compared to normal mucosa. *P*-values were calculated using DESeq2. (**B**) Cox hazard analysis of *RET* expression and survival in the TCGA HNSC and CESC cohort. (**C**) HNSC-associated *RET* mutations are plotted as a function of CADD score and compared to known oncogenic mutations. CLD, cadherin-like domain. Cys, cysteine-rich domain. TK, tyrosine kinase.

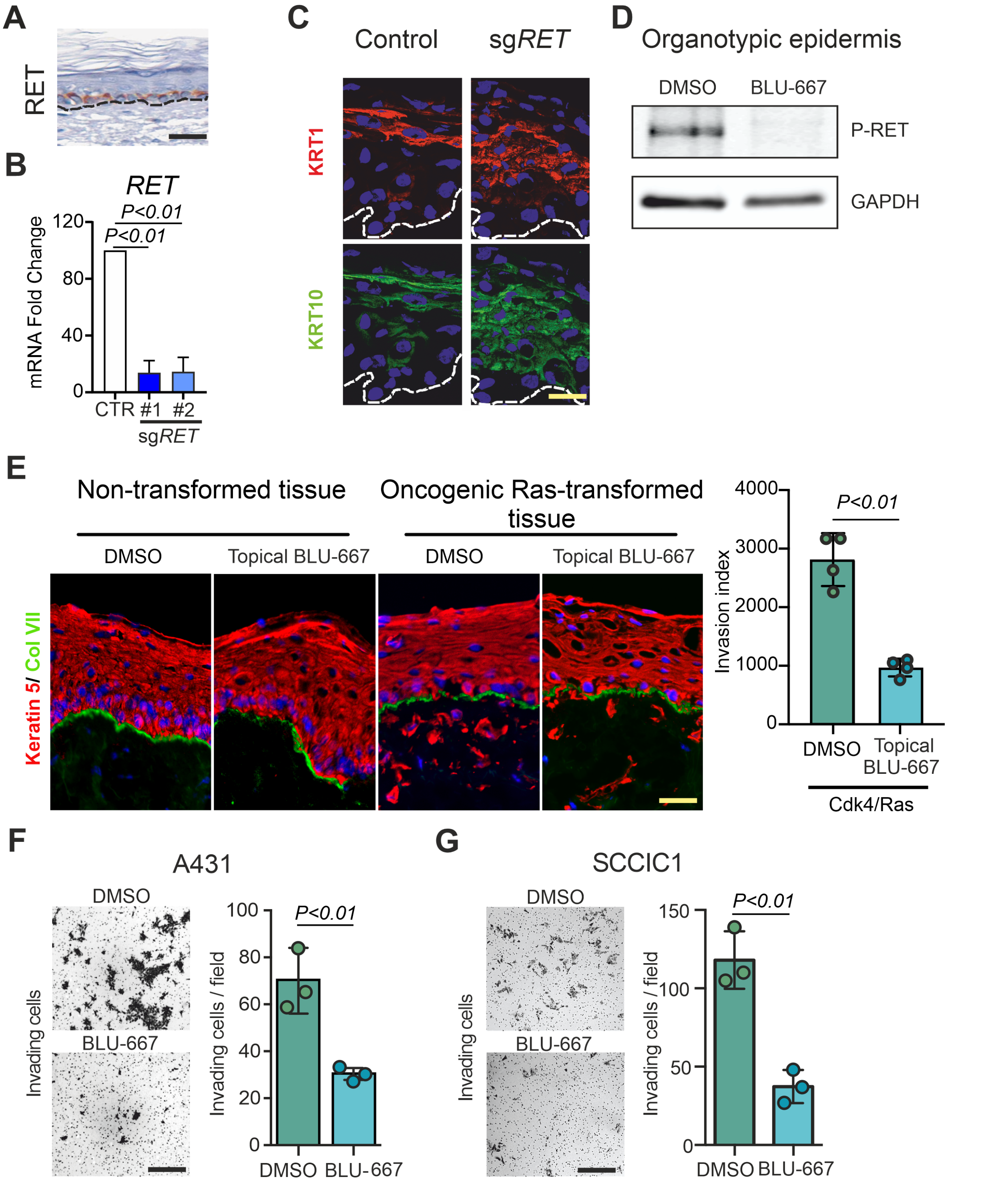


**Fig. S5. RET maintains epidermal progenitor cells in an undifferentiated state and its inhibition suppresses tumor-enabling behaviors in cSCC.** (**A**) RET expression in normal human skin available from v20.proteinatlas.org (https://www.proteinatlas.org/ENSG00000165731-RET/tissue/skin#img) (**B**) Quantitation of *RET* mRNA expression in primary human keratinocytes treated with two independent sgRNAs (sg*RET*) compared to a non-targeting control. Bars represent the mean of three technical replicates ± s.d. and similar results were confirmed in two experiments. (**C**) Differentiation proteins detected by immunofluorescence in red (keratin 1) and green (keratin 10), dotted line indicates basement membrane zone. (**D**) Western blot analysis of RET and phospho-RET expression in human skin organoids grown for seven days on dermis in the presence or absence of BLU-667 (15 nM). Data are representative of two experiments. (**E**) Keratin 5 (red) and collagen VII (green) immunofluorescence of oncogenic Ras-transformed human skin organoids after seven days of growth on dermis and in the presence or absence of topical BLU-667. Data are representative of three replicates. Quantitation of invasion index using four fields of view. The invasion index is calculated from the integrated density of keratin 5 staining beneath the basement membrane divided by the length of the basement membrane. (**F**,**G**) Quantitation of Matrigel invasion assays in human cSCC cell lines. For each cell line, three fields of view were used for quantitation and data are representative of three independent experiments. Bars represent the mean ± s.d. and significance was calculated using a two-tailed t-test. All scale bars, 100 µm.

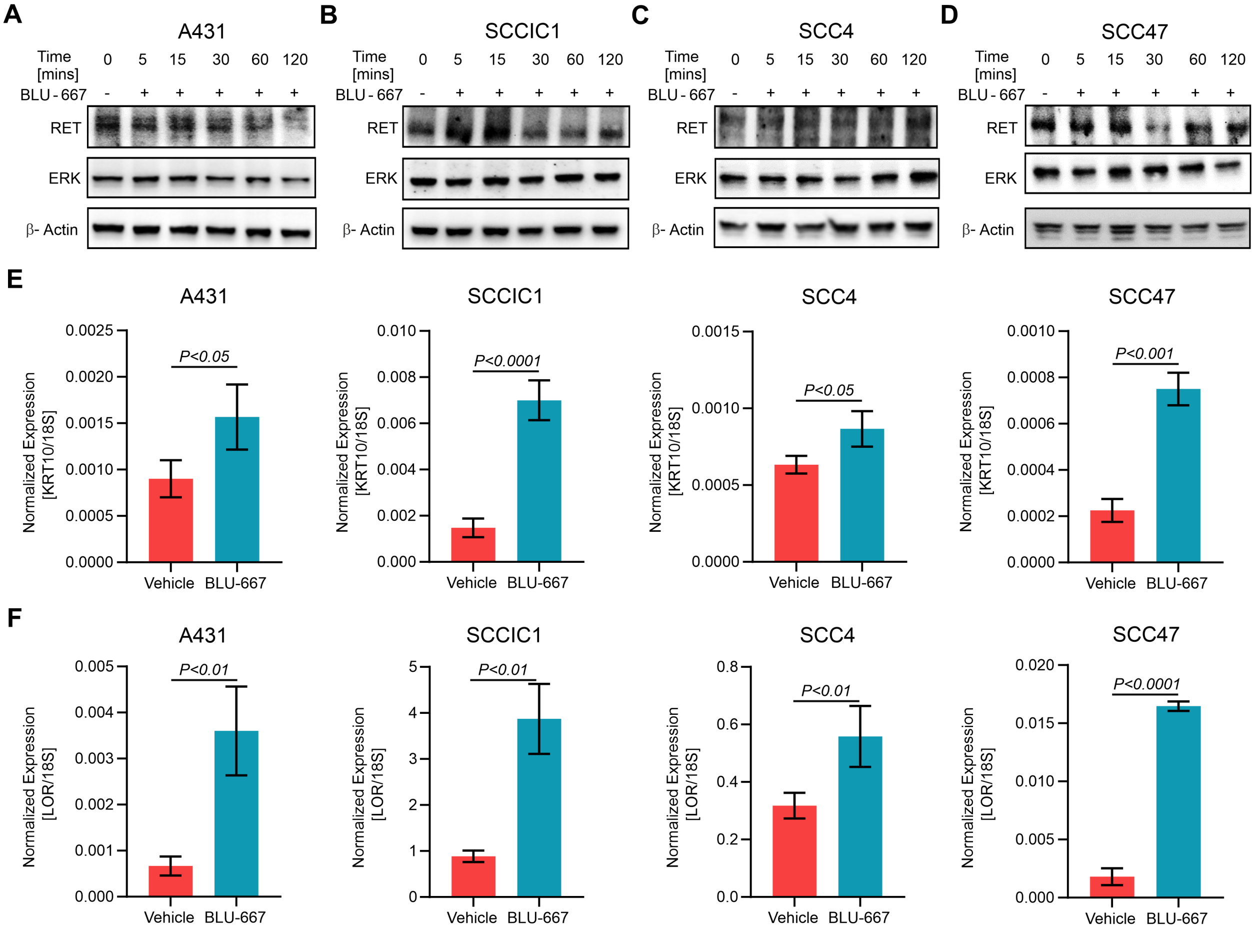


**Fig. S6. RET inhibition induces differentiation in SCC spheroids.** Western blot analysis of RET and ERK in A431 (**A**), SCCIC1 (**B**), SCC4 (**C**), and SCC47 (**C**) cells treated with BLU-667. The expression of differentiation markers keratin-10 (**E**) and loricrin (**F**) increases in SCC cell lines treated with BLU-667 (15 nM) for 24 hours compared to DMSO. Data were obtained by qRT-PCR performed in triplicate and are representative of three independent experiments.

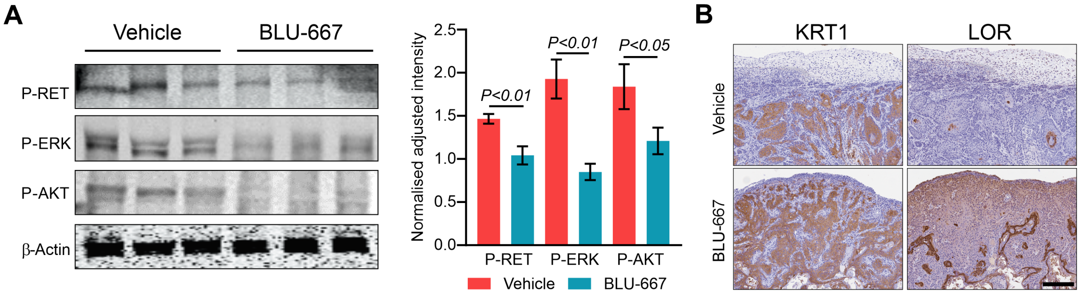


**Fig. S7. BLU-667 inhibits RET signaling and enhances differentiation in vivo.** (**A**) Immunoblotting of tumor lysates from A431 xenografts treated with BLU-667 or vehicle (n=3 tumors per treatment group). Quantitation of densitometry; bars represent mean $\pm$ s.d. and *P* values were determined using a two-tailed t-test. (**B**) Keratin-1 and loricrin immunohistochemistry of representative A431 xenografts treated with BLU-667 or vehicle. Scale bar, 100 μm.

**Supplementary Table 1. Univariable relationship between gene expression and overall survival in TCGA cervical (CESC) and head and neck cancer (HNSC).** HR, hazard ratio. CI, confidence interval.

| Gene | HR | P-value | 95% CI |
|---|---|---|---|
| *MAB21L4* | 0.942 | 0.0161 | 0.897-0.989 |
| *RET* | 1.115 | 0.000110 | 1.055-1.178 |

**Supplementary Table 2. Primer Sequences**

| Gene | Forward Sequence | Reverse Sequence |
|---|---|---|
| *RPL32* | AGGCATTGACAACAGGGTTC | GTTGCACATCAGCAGCACTT |
| *KRT1* | GAAGTCTCGAGAAAGGGAGCA | ATGGGTTCTAGTGGAGGTATCTA |
| *KRT10* | GCAAATTGAGAGCCTGACTG | CAGTGGACACATTTCGAAGG |
| *LOR* | CTCTGTCTGCGGCTACTCTG | CACGAGGTCTGAGTGACCTG |
| *CALML5* | CTACGAGGAGTTCGCGAGG | GTTTCCCATCCACCGACCAG |
| *CASP14* | GGCCTGTCGAGGAGAACAAA | ACGTGCAAGGCATCTGTGTA |
| *LCE3D* | GCTGCTTCCTGAACCAC | GGGAACTCATGCATCAAG |
| *TGM1* | CTTCAAGAACCCCCTTCCCG | TGAGGATCTTGGGCCTCTGT |
| *MAB21L4* | TGGACCACCGAGGATACCTT | AGCGAACCTGGTGCGATG |
| *OVOL1* | GCTCCCGGCTTCAGTTA | GGGCTGTGGTGGGCAGAA |
| *CACYBP* | CCCATCTCTGTGGAAGGCAG | CTGGGTCAGGTAATCCCACC |
| *RET* | GGCTTGTCCCGAGATGTTTA | GCAGGACACCAAAAGACCAT |
| *GAPDH* | GAAGAGAGAGACCCTCACTGCTG | ACTGTGAGGAGGGGAGATTCAGT |
| *18S* | CGGCTACCACATCCAAGGAA | GCTGGAATTACCGCGGCT |
